# HECTD1 is both a positive regulator and substrate of caspase-3 activity during apoptotic cell death

**DOI:** 10.1101/2023.02.17.528998

**Authors:** Nico Scholz, Florian Siebzehnrubl, Julien D. F Licchesi

## Abstract

Programmed cell death is a complex and tightly regulated sequence of events that determines cell fate during tissue homeostasis, development, and pathogenesis. The small protein modifier ubiquitin mediates important regulatory functions during cell death by regulating the stability and activity of checkpoint proteins and the assembly of cell death signalling complexes. The caspase family of cysteine aspartases are essential effectors of apoptotic cell death. Components of the ubiquitin system including RING ubiquitin ligases XIAP, MDM2, RBX1; RBR E3 ubiquitin ligases Parkin and LUBAC; and HECT E3 ubiquitin ligases NEDD4 and Itch are also substrates of caspase-mediated cleavage. In the case of NEDD4 and Itch, the single cleavage event occurs outside of the catalytic HECT domain and it remains unclear whether such cleavage events impact on ubiquitin ligase activity and/or function. Here, we identified the E3 ubiquitin ligase HECTD1 as the third HECT E3 cleaved by caspase-mediated cleavage during apoptotic cell death, in a manner which does not affect the integrity of the catalytic C-ter HECT domain. We mapped the single cleavage event to DFLD^1664^↓S and showed that the cleaved C-ter product, which contains the HECT ligase domain, is as stable as the endogenous full length protein. We also found that HECTD1 transient depletion led to reduced caspase-3 activity, but not caspase 8 nor 9. Furthermore, we also identified caspase-3 as the protease responsible for HECTD1 cleavage at Asp1664 suggesting that HECTD1 and caspase-3 might be part of a novel feedback loop mechanism during apoptotic cell death. This study highlight novel crosstalk between cell death mechanisms and the ubiquitin system and raises important questions on whether proteolytic cleavage of E3 ubiquitin ligases might represent an underappreciated mode of regulation during cell death mechanisms.

## 1. Introduction

Cell death is a necessary feature of all living organisms, from shaping embryonic development to maintaining tissue homeostasis in mammals. Apoptosis is the main signalling cascade driving programmed cell death and in contrast to other cell death pathways including necroptosis and pyroptosis, it does so through a non-lytic mode of action and is further immunologically silent (1–3). During apoptosis auto-processing of initiator caspases facilitates signal integration of apoptotic stimuli and subsequent signal propagation through the apoptotic response network, culminating in coordinated cleavage of a pre-defined set of substrates by activate effector caspase (4). Extrinsic apoptosis is mediated by death receptor signalling pathways, namely TNF-α:TNFR1, TRAIL-DR4/5 and FasL-Fas, which converge on activation of caspase 8 thus selectively driving apoptotic cell death (5, 6). Although initially discovered in the context of apoptosis, the initiator caspase 8 is now recognised as a molecular switch to coordinate apoptosis, necroptosis, and pyroptosis (7).

Protein cleavage during programmed cell death extends beyond the processing of inactive zymogens into active caspases; for example, Poly (ADP-ribose) polymerase (PARP) is cleaved by caspase 7 (8, 9). PARP cleavage serves as a regulatory mechanism which is functional during apoptosis as evidenced by the caspase resistant PARP-1 knock-in mutant mouse strain which showed reduced intestinal and renal ischemia/reperfusion injury due to downregulation of NF-κB-mediated transcription of inflammatory genes (10). Similarly, caspase-resistant PARP-1 promotes necrosis and accelerates apoptosis during TNF-α and staurosporine induced cell death suggesting an active yet complex regulatory role (11–13) (Herceg and Wang 1999; Boulares et al. 1999; Los et al. 2002). Caspase 8 is essential for apoptosis but also to prevent the induction of death-receptors during necroptosis. Recent data revealed RIPK1 as target of caspase 8-mediated cleavage which explains how caspase 8 activity limits apoptosis and necroptosis (14). NEDD4-binding protein 1 (NBP4) was also identified as caspase 8 substrate and this mechanism prevents NBP4 inhibitory function of cytokine and chemokine responses (15).

Pattern recognition receptors (PRRs), including Toll-like receptors (TLRs) and Nod-like receptors (NLRs), recognise pathogen-and danger-associated molecular patterns (PAMPS and DAMPs) which leads to assembly of the inflammasome, a multiprotein oligomeric innate immune complex which also contains caspase 1 (16). Caspase 1 is a key component of this complex and its activation triggers the maturation of pro-inflammatory cytokines IL-1β and IL-18 via proteolytic processing (1). Pyroptosis can also be triggered by sensing of intracellular LPS from gram-negative bacteria and this induces assembly of the non-canonical inflammasome. As a result, caspases 4/5 mediate the cleavage and activation of the pore-forming protein Gasdermin D, the effector of pyroptosis (17).

Proteolytic cleavage by caspases has also been reported for a handful of E3 ubiquitin ligases: for RING E3 ligases MDM2 and IAPs XIAP (BIRC4) and cIAP1 (BIRC2) and ML-IAP (BIRC7), the Cullin-RING E3 ligase subunit RBX1, the RING-in between-RING E3 ligases HOIP/HOIL1 (LUBAC subunits) and Parkin, and HECT E3 ligases Itch and NEDD4 (18–28). Caspase-mediated cleavage can have different effects on E3 ubiquitin ligase activity and function. For example, caspase-3 removes the p53 binding site of MDM2 and this abrogates its ability to ubiquitinate p53; caspase 1 cleaves and inactivates Parkin ligase activity which contributes to dopaminergic neuronal cell death; caspase-3 and 6 cleave RNF31 at multiple sites preventing it from activating NF-κB signalling (18,20,29). In contrast, cleavage of the E3 ubiquitin ligase ML-IAP (BIRC7) by caspase-3 and 7, yields a protein with an intact Inhibitor of Apoptosis (IAP) repeat and a RING domain. As a result, cleaved ML-IAP (BIRC7) instead gains pro-apoptotic functions (25).

Homologous to E6AP C-Terminus (HECT) E3 ubiquitin ligases are also targeted by caspase cleavage. Nedd4 is cleaved at Asp^598^ by caspase 1, 3 and 7 (*in vitro* at least), which results in the removal of the N-terminal C2 domain and destabilisation of the WW-HECT cleavage product (19). Itch is cleaved at Asp^240^ by caspases 3, 6, and 7 (*in vitro* at least) thus also separating the N-terminal C2 domain from the C-terminal WW-HECT cleavage product; however, cleavage was shown to not impact on HECT E3 ligase catalytic activity (24). Therefore, Itch cleavage resulting in the removal of the N-terminal C2 domain is akin to Ca^2+^-mediated release of NEDD4 C2:HECT auto-inhibition and subsequent activation of NEDD4 E3 ubiquitin ligase activity. Indeed, deletion of the C2 domain resulted in the constitutive activation of NEDD4 ubiquitin ligase activity (*in vitro*) (30).

The HECT E3 ubiquitin ligase HECTD1 plays roles in a variety of cellular processes including base excision repair, cell migration, transcriptional regulation, ribosome biogenesis, cell cycle regulation and embryonic development (31–37). Other studies have also hinted at putative functions for HECTD1 in inflammation/cell death pathways. In a mouse model of colitis, deletion of the carboxypeptidase inhibitor Latexin (LTX) enhanced the interaction between HECTD1 and Rsp3 which contributed to IκB-α ubiquitylation and degradation, and thus an increased pro-inflammatory response (38). A putative role for HECTD1 in driving inflammation was also suggested in mouse astrocytes where LPS treatment was found to increase HECTD1 protein levels and to exacerbate LPS-induced neuroinflammation through the p38-JNK axis (39). In addition to pro-inflammatory roles, HECTD1 is also proposed to exhibit pro-apoptotic functions. Gene set enrichment analysis of HECTD1 knockdown T47D breast cancer cells revealed a decrease in apoptotic gene signatures (33). In addition, HECTD1 was shown to be responsible for Apollon (BIRC6) ubiquitylation and its subsequent degradation, thereby promoting cell death in triple negative breast cancer (40).

In this study we report the discovery that HECTD1 is cleaved upon induction of intrinsic or extrinsic apoptotic cell death, and we identified caspase-3 as the sole caspase responsible for HECTD1 cleavage. We mapped HECTD1 cleavage to the P1 residue Asp^1664^ in the caspase consensus motif DFLD which we found is conserved in vertebrates including zebrafish. We then carried out functional assays to explore HECTD1 function in the context of cell death and found that it is required for full onset of apoptotic cell death (or activation of caspase-3). Our data reveal a so far unreported role for HECTD1 as a novel nexus during apoptotic cell death which warrants further work to expand on the mechanism involved.

## 2. Materials and Methods

### 2.1. Mammalian cell culture reagents

**Table 1.** Cell lines. Cell line, source, storage, and media composition.

| Reagent | Source | Storage | Media composition |
| --- | --- | --- | --- |
| **HEK293T** | Purchased from<br>atcc.org | Liquid nitrogen | DMEM; 10% (v/v) FBS |
| **HeLa** | Purchased from<br>atcc.org | Liquid nitrogen | DMEM; 10% (v/v) FBS |
| **THP-1** | Contributed by<br>Professor Ruth<br>Massey's lab<br>(University of Bristol) | Liquid nitrogen | RPMI 1640; 2 mM L-glutamine, 10% (v/v) FBS |

**Table 2.**
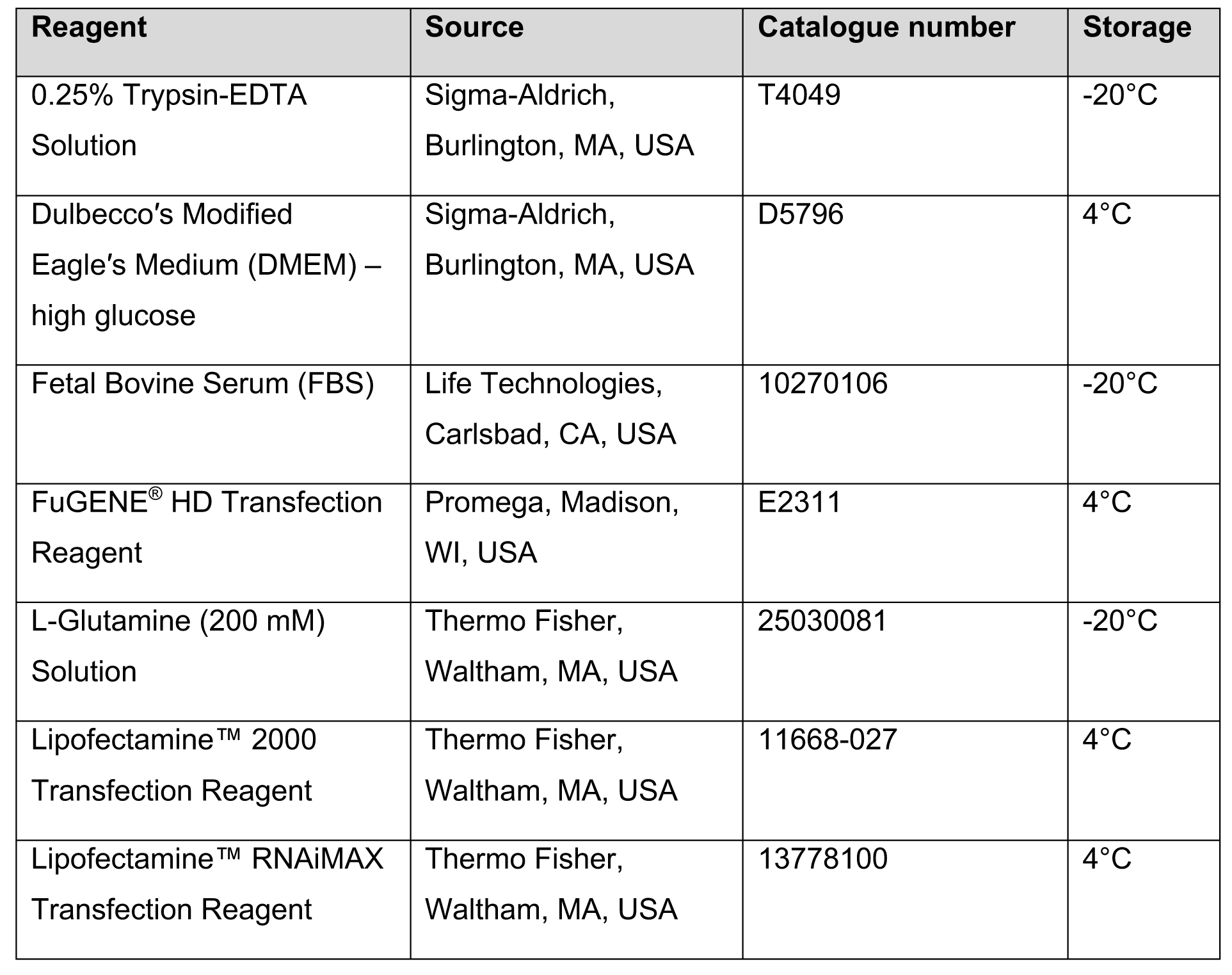
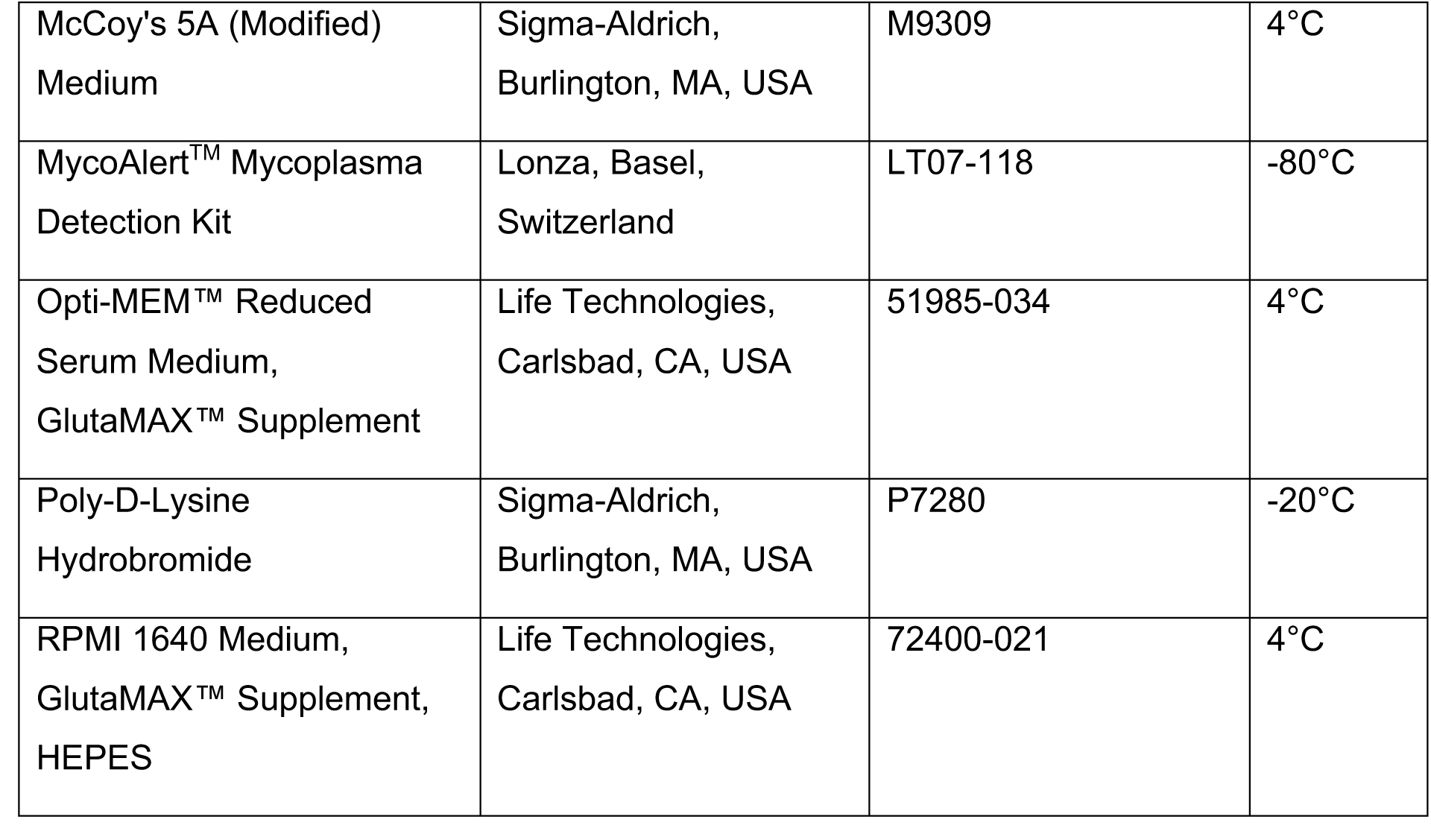
Mammalian cell culture reagents. Reagent, source, catalogue number, storage.

| Reagent | Source | Catalogue number | Storage |
| --- | --- | --- | --- |
| 0.25% Trypsin-EDTA<br>Solution | Sigma-Aldrich,<br>Burlington, MA, USA | T4049 | -20°C |
| Dulbecco's Modified<br>Eagle's Medium (DMEM) –<br>high glucose | Sigma-Aldrich,<br>Burlington, MA, USA | D5796 | 4°C |
| Fetal Bovine Serum (FBS) | Life Technologies,<br>Carlsbad, CA, USA | 10270106 | -20°C |
| FuGENE<sup>®</sup> HD Transfection<br>Reagent | Promega, Madison,<br>WI, USA | E2311 | 4°C |
| L-Glutamine (200 mM)<br>Solution | Thermo Fisher,<br>Waltham, MA, USA | 25030081 | -20°C |
| Lipofectamine<sup>™</sup> 2000<br>Transfection Reagent | Thermo Fisher,<br>Waltham, MA, USA | 11668-027 | 4°C |
| Lipofectamine<sup>™</sup> RNAiMAX<br>Transfection Reagent | Thermo Fisher,<br>Waltham, MA, USA | 13778100 | 4°C |
| McCoy's 5A (Modified)<br>Medium | Sigma-Aldrich,<br>Burlington, MA, USA | M9309 | 4°C |
| MycoAlert™ Mycoplasma<br>Detection Kit | Lonza, Basel,<br>Switzerland | LT07-118 | -80°C |
| Opti-MEM™ Reduced<br>Serum Medium,<br>GlutaMAX™ Supplement | Life Technologies,<br>Carlsbad, CA, USA | 51985-034 | 4°C |
| Poly-D-Lysine<br>Hydrobromide | Sigma-Aldrich,<br>Burlington, MA, USA | P7280 | -20°C |
| RPMI 1640 Medium,<br>GlutaMAX™ Supplement,<br>HEPES | Life Technologies,<br>Carlsbad, CA, USA | 72400-021 | 4°C |

### 2.2. Biochemical reagents

**Table 3.**
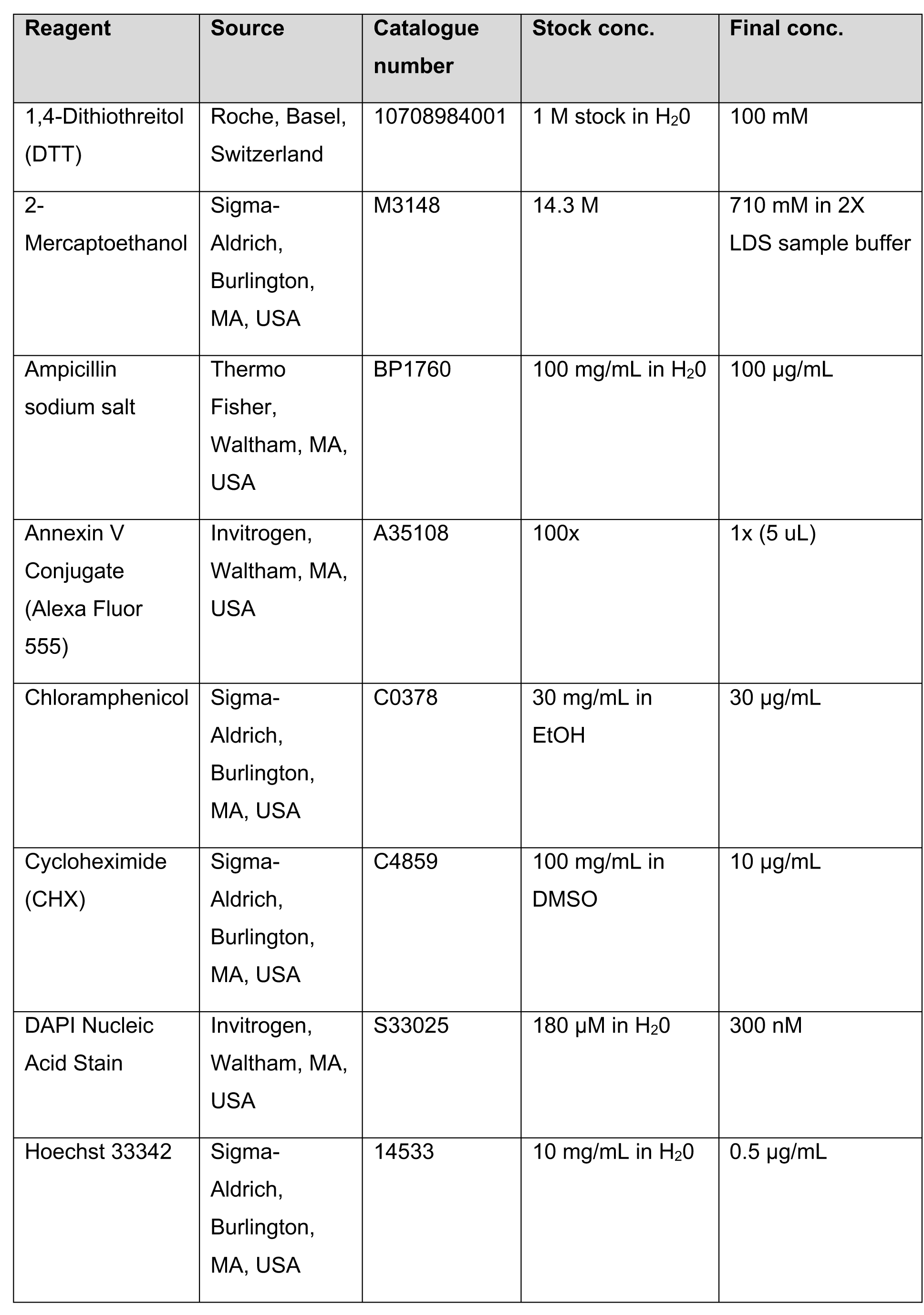
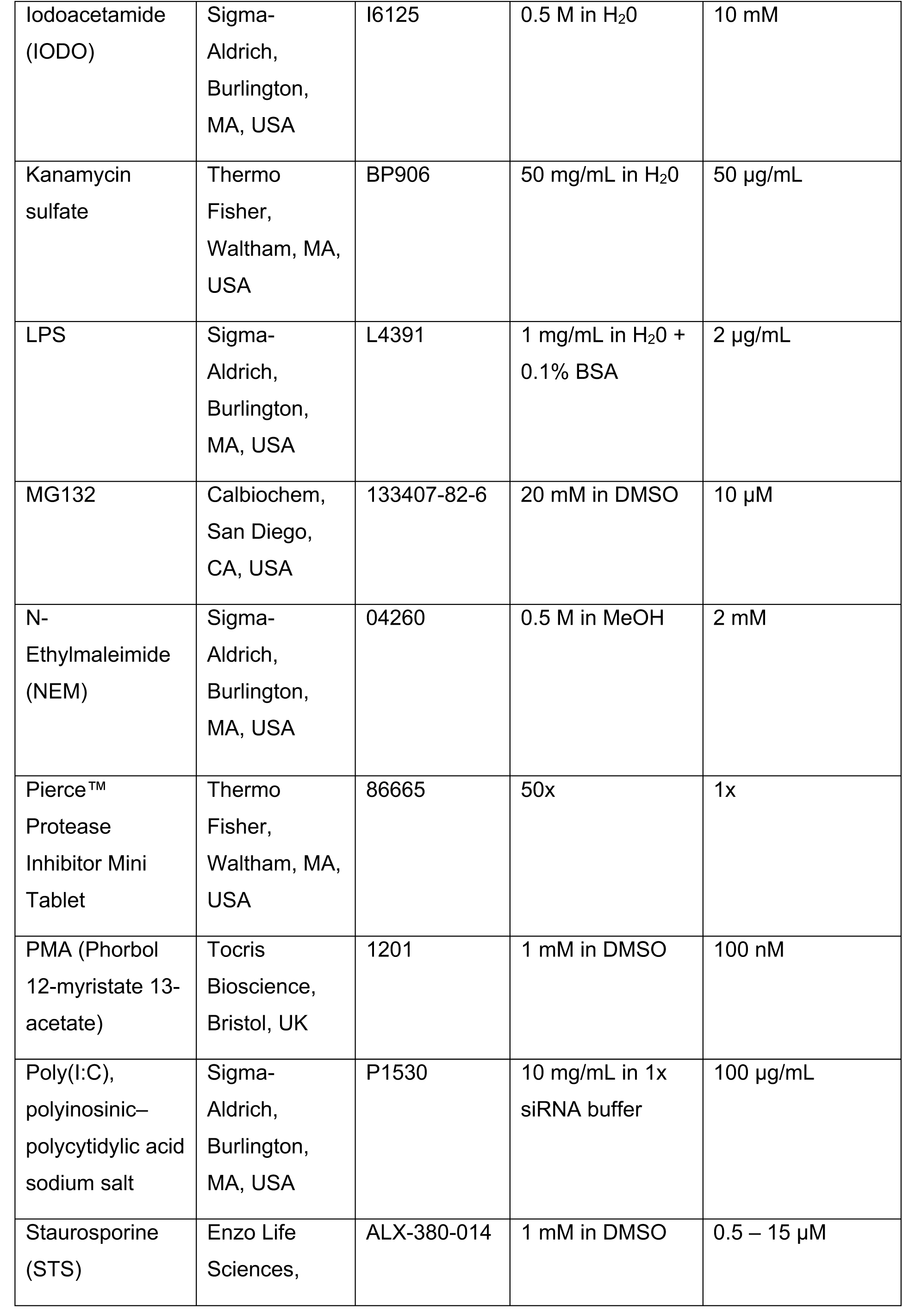
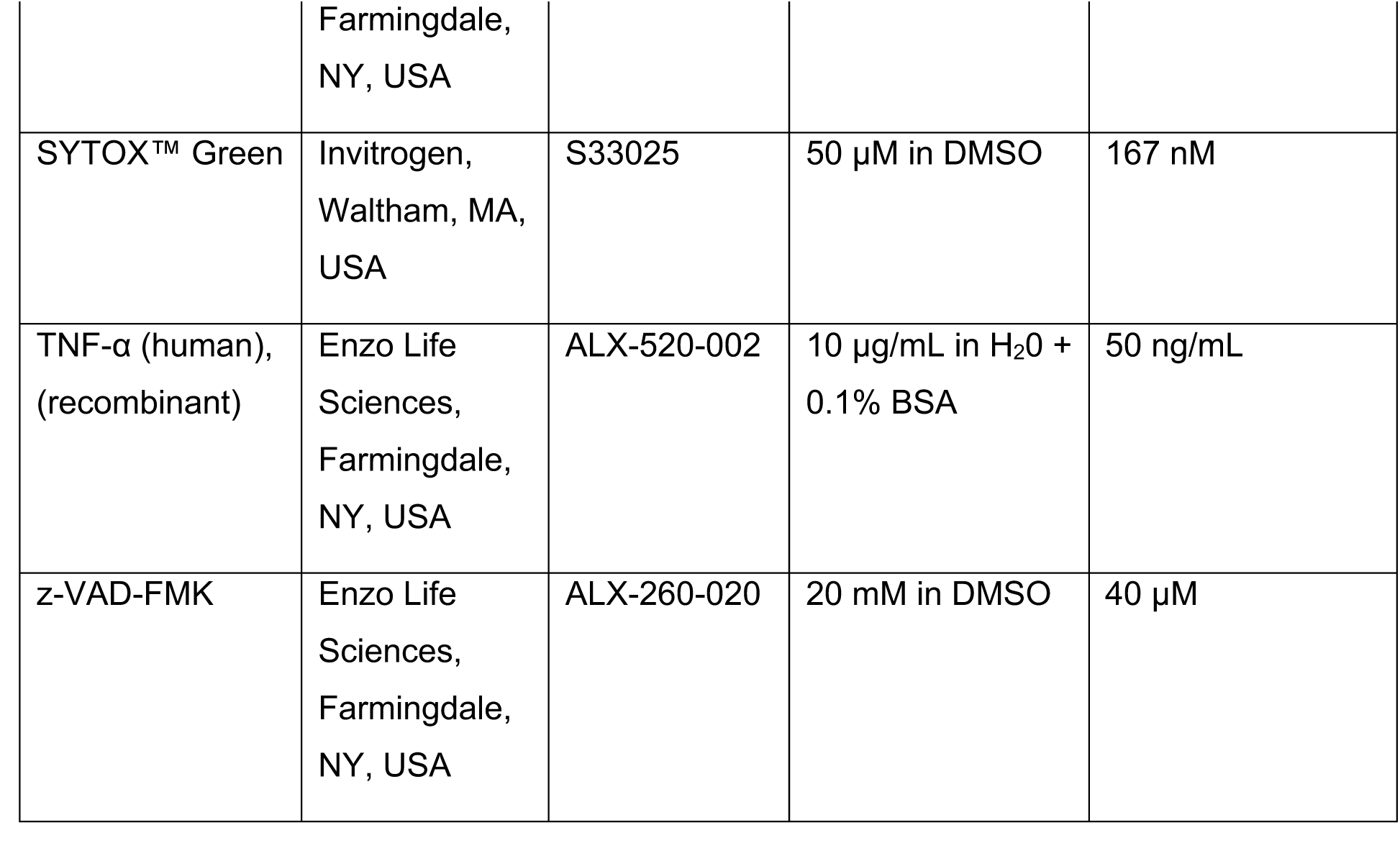
Biochemical reagents. Reagent, Source, catalogue number, [stock], and [final].

**Table 4.** Kits. Kits, source, and catalogue number.

| Kits | Source | Catalogue number |
| --- | --- | --- |
| Anti-FLAG® M2 Magnetic Beads | Sigma-Aldrich,<br>Burlington, MA, USA | M8823 |
| Bolt™ 4 to 12%, Bis-Tris, 1.0 mm,<br>Mini Protein Gel, 15-well/17-well | Invitrogen, Waltham,<br>MA, USA | NW04125BOX/<br>NW04127BOX |
| Caspase-3, Caspase-8 and<br>Caspase-9 Multiplex Activity Assay<br>Kit (Fluorometric) | Abcam, Cambridge,<br>UK | ab219915 |
| Dual-Luciferase® Reporter Assay<br>System | Promega, Madison,<br>WI, USA | E1910 |
| ECL™ Prime Western Blotting<br>System | GE Healthcare,<br>Chicago, IL, USA | RPN2232 |
| GeneJET Gel Extraction Kit | Thermo Fisher,<br>Waltham, MA, USA | K0691 |
| GeneJET Plasmid Maxiprep Kit | Thermo Fisher,<br>Waltham, MA, USA | K0491 |
| GeneJET Plasmid Miniprep Kit | Thermo Fisher,<br>Waltham, MA, USA | K0503 |
| HiSpeed Plasmid Midi Kit | Qiagen, Hilden,<br>Germany | 12643 |
| Pierce™ BCA Protein Assay Kit | Thermo Fisher,<br>Waltham, MA, USA | 23225 |
| QIAprep Spin Miniprep Kit | Qiagen, Hilden,<br>Germany | 27106 |
| QIAquick Gel Extraction Kit | Qiagen, Hilden,<br>Germany | 28704 |
| Trans-Blot Turbo RTA Midi 0.2 µm<br>PVDF Transfer Kit | Bio-Rad, Hercules,<br>CA, USA | 170-4273 |

### 2.3. Cloning Reagents

**Table 5.** Cloning reagents. Reagents, source, and catalogue number.

| Reagent | Source | Catalogue number |
| --- | --- | --- |
| 1 Kb Plus DNA Ladder | Invitrogen, Waltham, MA, USA | 10787018 |
| FastDigest Restriction Enzymes | Thermo Fisher, Waltham, MA, USA | Various |
| GelStar™ Nucleic Acid Gel Stain | Lonza, Basel, Switzerland | 50535 |
| In-Fusion® HD Cloning Kit | Takara Bio, Kusatsu, Shiga, Japan | 639649 |
| KOD Hot Start DNA Polymerase | Sigma-Aldrich, Burlington, MA, USA | 71086 |
| Monarch® PCR & DNA Cleanup Kit | New England Biolabs, Ipswich, MA, USA | T1030 |
| Phusion Hot Start II DNA polymerase | Thermo Fisher, Waltham, MA, USA | F549S |
| Stellar™ Competent Cells | Clontech Laboratories, Mountain View, CA, USA | 636763 |
| T4 DNA Ligase | New England Biolabs, Ipswich, MA, USA | M0202 |

### 2.4. Buffers and solution

**Table 6.** Buffers and solutions. Name, composition, and application.

| Name | Composition | Application |
| --- | --- | --- |
| 1X Annexin V Binding Buffer | Invitrogen, Waltham, MA, USA; V13241 | Flow Cytometry |
| 4X Bolt™ LDS Sample Buffer | Invitrogen, Waltham, MA, USA; B0007 | Western blot |
| 5X siRNA Buffer | Horizon Discovery, Waterbeach, UK; B-002000-UB-100 | RNA oligo resuspension |
| Coomassie Destaining Solution | 50% (v/v) MeOH, 10% (v/v) glacial acetic acid | Coomassie staining |
| Coomassie Staining Solution | 0.05% (w/v) Coomassie Brilliant Blue R-250, 50% (v/v) MeOH, 10% (v/v) glacial acetic acid | Coomassie staining |
| DNA Loading Dye Buffer (10X) | 40% (v/v) glycerol, 0.25% (w/v) bromophenol blue | Electrophoresis |
| GST Pulldown Buffer | 25 mM Tris pH 7.4, 150 mM NaCl, 5 mM DTT and 0.1% (v/v) NP-40 | Pulldown |
| Milk Blocking Buffer | 5% (w/v) oxoid skim milk in PBST | Western blot |
| NuPAGE™ MOPS SDS Running Buffer (20X) | Thermo Fisher, Waltham, MA, USA; NP0001 | SDS-PAGE |
| PBST | 1X PBS, 0.1% (v/v) Tween 20 | Western blot |
| Pierce™ Protein-Free T20 (PBS) Blocking Buffer | Thermo Fisher, Waltham, MA, USA; 37573 | Western blot |
| Recombinant Caspase Reaction Buffer | 50 mM Hepes, pH 7.2, 50 mM NaCl, 0.1 % CHAPS, 10 mM EDTA, 5 % Glycerol, and 10 mM DTT | Recombinant Caspase Assay |
| RIPA Lysis Buffer | 50 mM Tris pH 7.4, 150 mM NaCl, 2 mM EDTA, 1% (v/v) NP-40, 0.1% SDS | Western blot |
| SDS-PAGE Electrophoresis Running Buffer | 25 mM Tris, 192 mM glycine, 0.1% SDS, pH 8.3 | SDS-PAGE |
| TAE DNA Electrophoresis Buffer (50X) | 2 M Tris, 50 mM EDTA | Electrophoresis |
| Triton-X 100 Lysis Buffer | 50 mM Tris pH 7.5, 200 mM NaCl, 1% (v/v) Triton-X 100 | Pulldown |
| WB Sample Buffer | 2X LDS/100mM DTT | Western blot |
| Wet Transfer Buffer | 25 mM Tris pH 8.3, 192 mM glycine, 0.1% SDS, 20% (v/v) MeOH | Western blot |

### 2.5. Antbodies

### Primary Antibodies

**Table 7.**
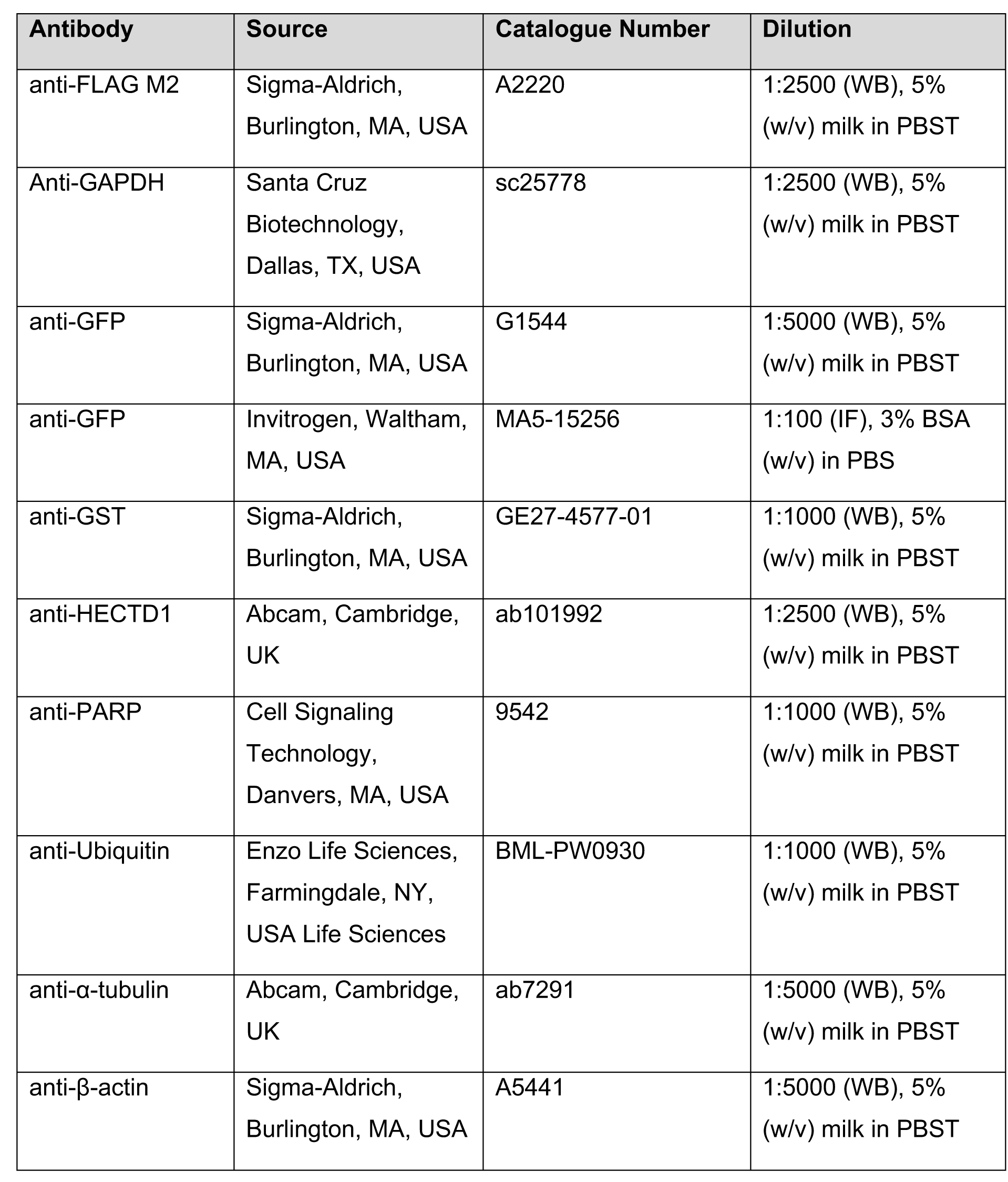
Primary antibodies. Antibody, source, catalogue number, and dilution.

| Antibody | Source | Catalogue Number | Dilution |
| --- | --- | --- | --- |
| anti-FLAG M2 | Sigma-Aldrich,<br>Burlington, MA, USA | A2220 | 1:2500 (WB), 5%<br>(w/v) milk in PBST |
| Anti-GAPDH | Santa Cruz<br>Biotechnology,<br>Dallas, TX, USA | sc25778 | 1:2500 (WB), 5%<br>(w/v) milk in PBST |
| anti-GFP | Sigma-Aldrich,<br>Burlington, MA, USA | G1544 | 1:5000 (WB), 5%<br>(w/v) milk in PBST |
| anti-GFP | Invitrogen, Waltham,<br>MA, USA | MA5-15256 | 1:100 (IF), 3% BSA<br>(w/v) in PBS |
| anti-GST | Sigma-Aldrich,<br>Burlington, MA, USA | GE27-4577-01 | 1:1000 (WB), 5%<br>(w/v) milk in PBST |
| anti-HECTD1 | Abcam, Cambridge,<br>UK | ab101992 | 1:2500 (WB), 5%<br>(w/v) milk in PBST |
| anti-PARP | Cell Signaling<br>Technology,<br>Danvers, MA, USA | 9542 | 1:1000 (WB), 5%<br>(w/v) milk in PBST |
| anti-Ubiquitin | Enzo Life Sciences,<br>Farmingdale, NY,<br>USA Life Sciences | BML-PW0930 | 1:1000 (WB), 5%<br>(w/v) milk in PBST |
| anti-α-tubulin | Abcam, Cambridge,<br>UK | ab7291 | 1:5000 (WB), 5%<br>(w/v) milk in PBST |
| anti-β-actin | Sigma-Aldrich,<br>Burlington, MA, USA | A5441 | 1:5000 (WB), 5%<br>(w/v) milk in PBST |

### Secondary antibodies

**Table 8.** Secondary antibodies. Antibody, source, catalogue number, and dilution.

| Antibody | Source | Catalogue number | Dilution |
| --- | --- | --- | --- |
| anti-Goat IgG<br>Peroxidase<br>Conjugated | Thermo Fisher,<br>Waltham, MA, USA | 31402 | 1:5000 (WB), 5%<br>(w/v) milk in PBST |
| anti-Mouse IgG<br>IRDye 680RD | LI-COR Biosciences,<br>Lincoln, NE, USA | 925-68070 | 1:10000 (IF), 5%<br>(w/v) milk in PBST |
| anti-Mouse IgG<br>Peroxidase<br>Conjugated | Sigma-Aldrich,<br>Burlington, MA, USA | AP124P | 1:5000 (WB), 5%<br>(w/v) milk in PBST |
| anti-Rabbit IgG<br>Peroxidase<br>Conjugated | Sigma-Aldrich,<br>Burlington, MA, USA | AP132P | 1:5000 (WB), 3%<br>BSA (w/v) in PBS |

## 3. Methods

### 3.1. Cloning

All primers were synthesised using the Sigma-Aldrich (Burlington, MA, USA) Custom DNA Oligo service and upon receipt resuspended in sterile water at 100 μM stock and 10 μM working concentration. Primers for mutagenesis were resuspended at a stock concentration of 1,000 pmol/μl and a working stock of 100 pmol/μl. All primer sequences are available upon request. All sequencing was carried out by Eurofins Scientific (Luxembourg, Luxembourg) TubeSeq sequencing service.

For restriction-based cloning, destination vectors (1 µg) were linearised using appropriate FastDigest restriction enzymes (Thermo Fisher, Waltham, MA, USA) for 1 h at 37°C. Linearised vectors were run on a 1% agarose gel and subsequently gel purified using the QIAquick Gel Extraction Kit (Qiagen, Hilden, Germany). Inserts were PCR amplified using either KOD Hot Start DNA Polymerase (0.5 U; ≤ 8 kb; Sigma-Aldrich, Burlington, MA, USA) or Phusion Hot Start II DNA polymerase (0.4 U; >8 kb; Thermo Fisher, Waltham, MA, USA) according to manufacturer’s instructions. Amplified inserts were confirmed on a 1-2% agarose gel and also gel purified using the QIAquick Gel Extraction Kit (Qiagen, Hilden, Germany). The isolated insert was the digested using appropriate restriction enzymes (Thermo Fisher, Waltham, MA, USA) for 1 h at 37°C and purified using the Monarch^®^ PCR & DNA Cleanup Kit (New England Biolabs, Ipswich, MA, USA). The ligation reaction was set up with T4 DNA Ligase (400 U; New England Biolabs, Ipswich, MA, USA) using a 3:1 molar ratio of insert:vector (50 ng) at 16°C O/N. Ligated plasmids were transformed into Stellar chemically competent cells (Clontech Laboratories, Mountain View, CA, USA) and subsequently sequence confirmed.

### 3.2. Site-directed Mutagenesis

Site directed mutagenesis was carried out to mutate individual candidate caspase cleavage sites. Mutagenesis primers were designed with the QuikChange Primer Design Tool (Agilent, Santa Clara, CA, USA). Plasmids were PCR amplified using either KOD Hot Start DNA Polymerase (0.5 U; ≤ 8 kb; Sigma-Aldrich, Burlington, MA, USA) or Phusion Hot Start II DNA polymerase (0.4 U; >8 kb; Thermo Fisher, Waltham, MA, USA) according to manufacturer’s instructions. Subsequently, template DNA was digested using 5 units of DpnI (Thermo Fisher, Waltham, MA, USA) for 1.5 h at 37°C. Digested mutagenesis reactions were then transformed into Stellar chemically competent cells (Clontech Laboratories, Mountain View, CA, USA) and subsequently sequence confirmed.

### 3.3. Domain-swapping Mutagenesis

Domain swapping was used to fuse Ubiquitin at the N-terminus of the C-ter HECTD1 product released by caspase cleavage in order to test its stability. Primers were designed with 20-nt homology arms on either side of the sequence to be amplified. Domains were PCR amplified using KOD Hot Start DNA Polymerase (0.5 U; Sigma-Aldrich, Burlington, MA, USA) according to manufacturer’s instructions. Amplified domains were confirmed on a 1-2% agarose gel and gel purified using the QIAquick Gel Extraction Kit (Qiagen, Hilden, Germany). The domain swapping PCR, using Phusion Hot Start II DNA polymerase (0.4 U; Thermo Fisher, Waltham, MA, USA), was set up with a 7:1 ratio of megaprimer (350 ng):plasmid backbone (50 ng) and an extended annealing time of 1 min. Reactions were then handled as for mutagenized reaction (detailed above).

**In-Fusion^®^ HD Cloning** was done according to manufacturer’s instructions, but briefly: Primers were designed using the Takara Bio primer design tool (Takara Bio, Kusatsu, Shiga, Japan). Inserts were PCR amplified using either KOD Hot Start DNA Polymerase (0.5 U; ≤ 8 kb; Sigma-Aldrich, Burlington, MA, USA) or Phusion Hot Start II DNA polymerase (0.4 U; >8 kb; Thermo Fisher, Waltham, MA, USA) according to manufacturer’s instructions. PCR reactions were cleaned up using the In-Fusion Cloning Enhancer. Linearised vectors (1 µg) were generated using appropriate FastDigest restriction enzymes (Thermo Fisher, Waltham, MA, USA) for 1 h at 37°C. Linearised vectors were run on a 1% agarose gel and subsequently gel purified using the QIAquick Gel Extraction Kit (Qiagen, Hilden, Germany). The In-Fusion cloning reaction was set up as calculated by the Takara In-Fusion molar ratio calculator. Sequences were validated by diagnostic restriction digest and sequencing.

### 3.4. Mammalian Cell Culture

HeLa cells, Human embryonic kidney 293T (HEK293T) cells and HEK293T were purchased from ATCC and cultured in DMEM (Sigma-Aldrich, Burlington, MA, USA) supplemented with 10% (v/v) Fetal Bovine Serum (Heat-inactivated FBS; Life Technologies, Carlsbad, CA, USA). HCT116 cells were cultured in McCoy′s 5A medium (Sigma-Aldrich, Burlington, MA, USA) supplemented with 10% (v/v) FBS (Life Technologies, Carlsbad, CA, USA). THP-1 cells (Gift from Ruth Massey’s laboratory, University of Bristol, Bristol, UK) were cultured in RPMI 1640 media (Life Technologies, Carlsbad, CA, USA) supplemented with 2 mM L-glutamine (Thermo Fisher, Waltham, MA, USA), 10% (v/v) FBS (Life Technologies, Carlsbad, CA, USA). All cells were at cultured 37°C and 5% CO2 in a humidified incubator. For differentiation into THP-1 macrophage-like cells, cells were seeded at 4×10^5^ cells/well in a 12-well plate and incubated with 100 nM phorbol 12-myristate 13-acetate (PMA; Tocris Bioscience, Bristol, UK) for 72 h. Media was changed after 72 h and cells were allowed to recover for 24 h (41). Confluent cells were sub-cultured by first washing cells with PBS and then detaching cells using 0.25% trypsin-EDTA (Sigma-Aldrich, Burlington, MA, USA). Cell suspensions were centrifuged for 1 min at 800 g (HeLa, HEK293T) or 5 min at 200 g (THP-1). Cell pellets were resuspended in fresh, complete media. All cell lines were tested for Mycoplasma contamination using the MycoAlert^TM^ Mycoplasma Detection Kit (Lonza, Basel, Switzerland) according to manufacturer instructions.

### 3.5. Transfection

DNA constructs were transfected 24 h post-seeding with Lipofectamine™ 2000 (Thermo Fisher, Waltham, MA, USA; HEK293T, HCT116) or FuGENE® HD (Promega, Madison, WI, USA; HeLa) according to manufacturer instructions using a 3:1 ratio. For example, 96-well plate: 50 ng DNA; 12-well plate: 500 ng DNA; 6-well plate: 1250 ng DNA. Unless otherwise indicated, assays were performed 24 h post-transfection. siRNA was transfected 24 h post-seeding with Lipofectamine™ RNAiMAX (Thermo Fisher, Waltham, MA, USA) according to manufacturer instructions using a 3:1 ratio. For example, 96-well plate: 1.2 pmol siRNA; 12-well plate: 12 pmol siRNA; 6-well plate: 30 pmol siRNA. Unless otherwise indicated, assays were performed 48 h post-transfection following a media change at 24 h post-transfection. siRNA (Dharamacon^TM^) were obtained from Horizon Discovery (Waterbeach, UK): On-Target Plus Non-Targeting Pool siRNA (D-001810-10-05), On-Target Plus *HECTD1* SMARTPool siRNA (J-007188-00), On-Target Plus *HECTD1* Individual #06 siRNA (J-007188-06), On-Target Plus *HECTD1* Individual #08 siRNA (J-007188-08).

### 3.6. Induction of cell death

The intrinsic mitochondrial apoptotic pathway was triggered by treating cells with staurosporine (STS; Enzo Life Sciences, Farmingdale, NY, USA), while extrinsic apoptotic cell death was induced by combined treatment with 50 ng/ml of TNF-α (Enzo Life Sciences, Farmingdale, NY, USA) and 100 mg/mL cycloheximide (CHX; Sigma-Aldrich, Burlington, MA, USA), or Poly(I:C) (TLR3-mediated apoptosis; Sigma-Aldrich, Burlington, MA, USA) at indicated concentrations in complete media. To block caspase activation, cells were simultaneously (unless otherwise indicated) incubated with the pan-caspase inhibitor z-VAD-FMK (Enzo Life Sciences, Farmingdale, NY, USA).

Pyroptosis was induced by transfecting THP-1 cells with 2 μg/mL lipopolysaccharide (LPS; Sigma-Aldrich, Burlington, MA, USA) using 2.5 μL/mL Lipofectamine™ 2000 (Thermo Fisher, Waltham, MA, USA) as previously described. Induction of pyroptosis was monitored by treating cells with 167 nM Sytox-Green and subsequent imaging and analysis using the Countess 3 Automated Cell Counter prior to Western blot analysis (Thermo Fisher, Waltham, MA, USA).

### 3.7. Flow cytometry

HeLa cells were seeded at 150,000 cells/well into 6-well plates and transfected as previously described. Apoptosis was induced using either 50 ng/mL TNF-α + 10 ug/mL CHX or 0.5 μM STS for indicated time points. Cells, including media, were harvested using 0.25 % trypsin-EDTA. To allow cell recovery, cells were re-suspended in 250 μL complete media and incubated for 30 min at 37°C and 5% CO2 in a humidified incubator whilst being agitated every few minutes. Approximately 2×10^5^ cells/sample were resuspended in 100 μL 1X Annexin V binding buffer (Invitrogen, Waltham, MA, USA) containing 5 uL Annexin V - Alexa Fluor 555 conjugate (Invitrogen, Waltham, MA, USA) and 300 nM DAPI (Invitrogen, Waltham, MA, USA) and then incubated for 15 min at room temperature. Following incubation, 400 μL 1X Annexin V binding buffer were added, samples placed on ice and analysed using a BD FACSAria III immediately. Data were analysed using FCS Express 7 Research Edition v7.14.0020.

### 3.8. Protein stability

To assess protein stability (endogenous or exogenous) HeLa cells were treated with 10 µg/mL CHX or 10 µg/mL CHX + 10 µM MG132 (Calbiochem, San Diego, CA, USA) in complete or reduced serum (0.5%) media. At indicated timepoints, cells were harvested and analysed by Western blotting.

### 3.9. Luciferase reporter assay

The Dual-Luciferase^®^ Reporter Assay (Promega, Madison, WI, USA) was performed according to manufacturer’s instructions. Briefly, HeLa cells were seeded at 8,000 cells/well in a 96-well plate and transfected as mentioned above. After treatment, cells were lysed in 20 µL passive lysis buffer for 15 min at room temperature. In a white 96-well plate, 50 µL luciferase assay reagent II were added to the lysate. After reading the plate, 50 µL freshly prepared 1X Stop & Glo^®^ Reagent were added and the plate read again. Plate readings were performed using the Promega GloMax®-Multi Detection System (Madison, WI, USA).

### 3.10. Western blot analysis

Cells were lysed in RIPA buffer supplemented with Pierce^TM^ Protease Inhibitor Mini Tablets (Thermo Fisher, Waltham, MA, USA) for 20 min on ice. Cell lysates were cleared by centrifugation at 13,000 rpm at 4°C. Supernatant was prepared using WB sample buffer (2X LDS/100 mM DTT) and denatured at 95°C for 5 min. Samples, typically 10-20 µg determined via Pierce™ BCA Protein Assay Kit (Thermo Fisher, Waltham, MA, USA), were run on Bolt™ 4 to 12%, Bis-Tris, 1.0 mm, Mini Protein Gels (Thermo Fisher, Waltham, MA, USA) for 70 min at 120 V in 1X NuPAGE^®^ MOPS SDS Running Buffer (Thermo Fisher, Waltham, MA, USA).

For immunoblotting, samples were transferred onto PVDF Transfer Membrane, 0.45 µm (Thermo Fisher, Waltham, MA, USA) using the Bio-Rad Mini Trans-Blot^®^ Wet Transfer System (Bio-Rad, Hercules, CA, USA) for 60 min at 100 V in a wet transfer system. As alternative transfer system, the Trans-Blot Turbo RTA Midi 0.2 µm PVDF Transfer Kit (Bio-Rad, Hercules, CA, USA) was used. Membranes were blocked in milk blocking buffer (5% (w/v) oxoid skim milk in PBST) for 1 h at room temperature.

Primary antibodies were diluted in milk blocking buffer or Pierce™ Protein-Free T20 (PBS) Blocking Buffer (Thermo Fisher, Waltham, MA, USA) and incubated for 1 h at room temperature or overnight at 4°C. Secondary antibodies were diluted in milk blocking buffer and incubated for 1 h at room temperature. Membranes were incubated with ECL™ Prime Western Blotting System (GE Healthcare, Chicago, IL, USA) for 1 min. Chemiluminescence detection was carried out using the Fusion SL Chemiluminescence and Fluorescence Imager (Vilber Lourmat, Collégien, France) and if appropriate, protein content was quantified using ImageJ software (42).

For anti-ubiquitin blots, samples were transferred onto Amersham™ Protran^®^ Western blotting nitrocellulose membranes, 0.45 µm (Cytiva, Marlborough, MA, USA) which were subsequently boiled for 10 min in distilled H2O (43).

### 3.11. Protein expression and purification

GST pulldown was performed using GST, GST-TRABID1-200, GST-TRABID1-200 TY>>LV and GST-TAB2 expressed in *E. Coli* as previously described (44, 45). Briefly, constructs were transformed in BL21(DE3) RIL chemically competent cells. A single colony was then inoculated in LB supplemented with 30 µg/ml chloramphenicol and either 100 µg/ml ampicillin or 50 µg/ml of kanamycin, depending on the construct, and incubated overnight at 30°C in a shaking incubator (220 rpm). The following day, 10 ml of overnight culture was used to inoculate 1 L of LB supplemented with either ampicillin or kanamycin depending on the constructs, and 50 µM of ZnCl2, and grown up to OD600 = 0.6 UA. A final concentration of 500 µM ZnCl2 was used to induce expression which was left to proceed overnight in a shaking incubator at 18°C. Cells were then harvested, lysed, and protein purification was carried out as previously described (45).

### 3.12. GST pull down

GST pulldown using ubiquitin binding domain was carried out as previously described (44, 45). Briefly, GST-tagged bait proteins were conjugated to Pierce^TM^ Glutathione Magnetic Agarose Beads (Thermo Fisher, Waltham, MA, USA) for 1 h at room temperature (20 µg bait protein/10 µL slurry/condition) in GST pulldown buffer (25 mM Tris pH 7.4, 150 mM NaCl, 5 mM DTT and 0.1% (v/v) NP-40). Following treatment, cells were harvested (1x 10cm^3^ dish/condition) and lysed in 1 ml Triton-X 100 lysis buffer (50 mM Tris pH 7.4, 150 mM NaCl, 1% (v/v) Triton X-1. 100) supplemented with Pierce^TM^ Protease Inhibitor Mini Tablets (Thermo Fisher, Waltham, MA, USA), 10 mM IODO (Sigma-Aldrich, Burlington, MA, USA) and 2 mM NEM (Sigma-Aldrich, Burlington, MA, USA). Samples were lysed for 20 min on ice followed by 15 min centrifugation at 13,000 rpm at 4°C. 50 µL sample were collected as input while the remainder was split evenly according to number of bait proteins used. Samples were incubated with beads in GST pulldown buffer supplemented with 0.5 mg/mL BSA for 2 h at room temperature. Subsequently, beads were washed 5x with GST pulldown buffer before being resuspended in 30 µl WB sample buffer (2X LDS/100 mM DTT).

### 3.13. Caspase-3, Caspase-8 and Caspase-9 Multiplex Activity Assay Kit

The Caspase-3, Caspase-8 and Caspase-9 Multiplex Activity Assay Kit (Fluorometric) (Abcam, Cambridge, UK) was performed according to manufacturer’s instructions. Briefly, HeLa cells were seeded at 4,000 cells/well in a black-walled 96-well plate and transfected as previously above. On the day of the experiment, apoptosis was induced as mentioned above and following treatment, caspase-3/8/9 substrate was diluted in assay buffer and added to the cells. After 45 min of incubation at room temperature (protected from light), the plate was read using the Promega GloMax®-Multi Detection System (Madison, WI, USA).

### 3.14. Caspase screening for HECTD1 cleavage

To generate full-length HECTD1 and assess which caspase(s) is responsible for its cleavage, we produced full-length HECTD1 in mammalian cells. pCMV-3xFLAG-Full-length mouse HECTD1 wild type (WT) or cleavage resistant (D1664E) were transiently transfected in HEK293T cells as described above. Two 10 cm dishes were used per construct. 24 h post-transfection, cells were harvested and lysed in Triton-X 100 lysis buffer (50 mM Tris pH 7.4, 150 mM NaCl, 1% (v/v) Triton X-100) supplemented with Pierce^TM^ Protease Inhibitor Mini Tablets (Thermo Fisher, Waltham, MA, USA). Anti-FLAG® M2 Magnetic Beads (Sigma-Aldrich, Burlington, MA, USA) were used to capture 3xFLAG-HECTD1 proteins, as per the manufacturer recommendation and as done previously (45, 46). Captured 3xFLAG-HECTD1 proteins were aliquoted and resuspended in recombinant caspase reaction buffer (50 mM Hepes, pH 7.2, 50 mM NaCl, 0.1 % CHAPS, 10 mM EDTA, 5 % Glycerol, and 10 mM DTT) ready for the addition of recombinant caspases.

Lyophilised caspases from the Recombinant Active Human Caspases Set IV (RayBiotech, Norcross, GA, USA) were resuspended in PBS, 15% glycerol (0.5 U/uL). 1 U active caspase was used per reaction and samples were incubated for 1 h at 37°C prior to the addition of 2X LDS/100 mM DTT to terminate the rection. Samples were then analysed by immunoblotting as detailed above.

### 3.15. Bioinformatics analysis

The PROTOMAP database contains novel caspase substrates that were identified in staurosporine-stimulated Jurkat T cells and was here utilised to obtain existing cleavage data of HECTD1. Data from 2, 4, and 6 h time points were used to identify cleavage patterns (https://www.scripps.edu/cravatt/protomap/) (47). Prospective HECTD1 caspase cleavage sites were identified using both CaspDB and ELM databases (http://elm.eu.org/) (48, 49).

### 3.16. Sequence alignment

Sequence from HECTD1 was retrieved from UniProt for Homos Sapiens, Mus musculus (Mouse), Danio rerio (Zebrafish), Rattus rattus (Rat), Gallus gallus (Chicken), Xenopus tropicalis and Xenopus laevi (Frog), Ciona interstinalis (pacific transparent sea squirt), Drosophila melanogaster (Fruit fly), Caenorhabditis elegans, Sacharomyces cerevisiase (Baker’s yeast) (50). Sequences were imported into Jalview, and alignment was carried out using MSA (MUSCLE) alignment and the default Clustal protein colour scheme was used (51, 52).

### 3.17. Statistical analysis

Statistical analysis was performed using GraphPad Prism version 9.1.1. for Windows, GraphPad Software, San Diego, California USA, www.graphpad.com. Statistical tests used are indicated in the relevant figure legends, but generally: Two-tailed student’s *t*-test (2 sample groups), One-Way ANOVA with Dunnett’s post hoc test (3+ sample groups), or Two-Way ANOVA with Šídák’s multiple comparisons test (3+ sample groups, 2+ categorical variables). Common significance thresholds used include ****p<0.0001, ***p<0.001, **p<0.01, and *p<0.05.

## 4. Results

### 4.1. HECT E3 ligases cleaved by caspase-mediated cleavage

Proteolytic processing is a widespread mechanism which can have different effects of protein fate and function. Protein cleavage activates zymogens into active caspases, removes peptide sequences which determine the folding, retention time and stability of ER resident and client proteins, or activate the pore forming protein gasdermin D during pyroptosis. Protein cleavage also generates proteins with “neo” N-and C-termini and the newly exposed residues can impact on protein stability which is orchestrated by the N-end and C-end rule pathways, respectively (53). This may suggest that protein cleavage is an underappreciated mode of protein regulation, and that cleaved products can have the capacity to act as “neo-proteins” with new functions.

Proteomics efforts including positive/negative N-termini enrichment strategies (DegraBase/COFRADIC) or in-gel digestion of lysates (PROTOMAP) have facilitated global mapping of caspase substrates (47,54,55). Quantitative MS-based enzymology has highlighted remarkable substrate specificity and unique cleavage rates indicating a caspase-specific substrate hierarchy reflective of their respective biological function (56). E3 ubiquitin ligases can also be targeted by caspase-mediated protein cleavage, including RING E3 ligases MDM2, BIRC2 and 7, the Cullin-RING E3 ligase subunit RBX1, the RING-in between-RING E3 ligases HOIP/HOIL1 (LUBAC subunits) and Parkin. HECT E3 ligases including Itch and NEDD4 have also been identified as caspase substrates and interestingly this proteolytic event does not appear to affect ubiquitin ligase activity, at least in the case of Itch (19, 24).

Global mapping of the topography of proteolytic events using the PROTOMAP database has been a powerful resource to identify and study protein cleavage events. Indeed, the PROTOMAP database revealed that cleavage events were enriched for in inter-domain regions and, in addition, often resulted in stable cleavage products which could indicate functionality (47). PROTOMAP datasets were obtained from SDS-PAGE/1D reverse-phase LC-MS/MS analysis of Jurkat T cells treated with staurosporine (STS) for 2, 4 or 6 h. Physical separation of SDS-PAGE gels into fixed interval bands in combination with the detection of specific peptides by MS enabled the 3-dimensional mapping of cleavage events by molecular weight and protein sequence, respectively, and therefore providing information on the topography and magnitude of proteolytic events in apoptosis.

To try and explore the possibility that additional HECT E3 ubiquitin ligases might be subject to proteolytic processing by caspases during apoptosis, we explored the PROTOMAP database and identified 6 HECT E3s with evidence of caspase cleavage (47). Further analysis of this list revealed that Itch, NEDD4 and HECTD1 were the only HECT ligases with a peptide signature and banding pattern indicative of a single cleavage site, located upstream of the C-terminal HECT domain (**Figure 1A-C**). While NEDD4 and Itch have already been defined as caspase substrates, HECTD1 cleavage has so far not been explored and we therefore set out to validate and expand on this finding (19, 24).

**Figure 1.**
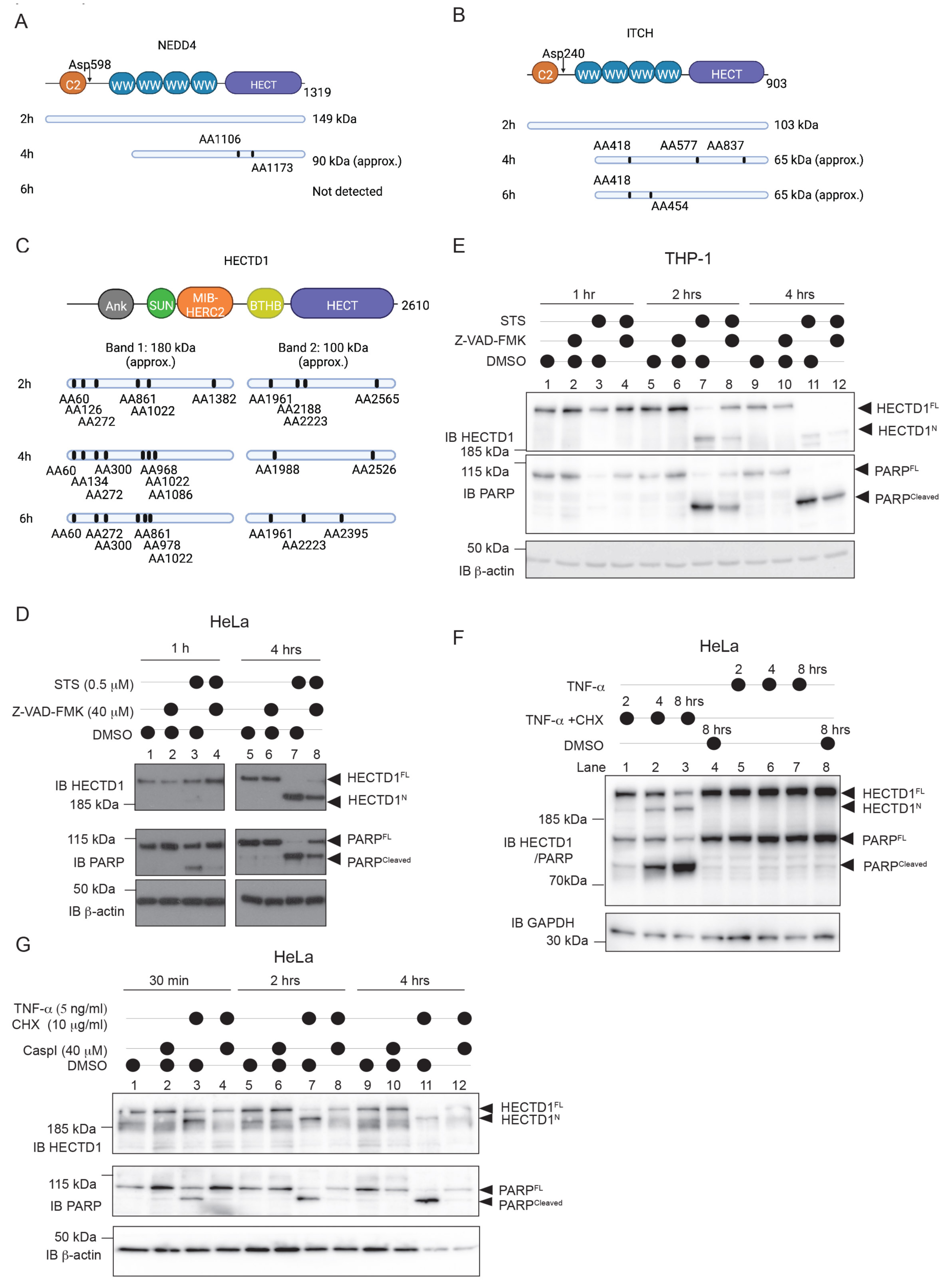
HECTD1 is proteolytically cleaved upon activation of intrinsic or extrinsic cell death pathways. **(A-C)** Pattern of PROTOMAP cleavage for NEDD4 **(A)**, Itch **(B)** and HECTD1 **(C)**. Top panels show the domain organisation of NEDD4, bottom panels show peptides identified following 2, 4 and 6 h of treatment of Jurkat T-cells with staurosporine (data from PROMOMAP). Cleavage of HECTD1 upon staurosporine treatment yielded two bands and based on the peptide identified, the cleavage site is likely located between the residue aa1382 and aa1961. **(D)** Induction of intrinsic apoptosis by STS treatment (1 μM) triggers HECTD1 cleavage in HeLa cells. The cleavage could be blocked by pre-treating cells with z-VAD-FMK (2 h pre-treatment at 40 μM, Lane 8 vs. 7). Cells were harvested at the indicated timepoints, lysed using RIPA buffer, and samples were resolved on a 4-12% Bis-Tris PAGE. Membranes were probed using HECTD1 antibody (Ab101992; note the epitope is within the first 50 residues). PARP was used as control to validate induction of apoptotic cell death and caspase activation and GAPDH was used as loading control. Bands corresponding to full-length and cleaved HECTD1 and PARP are indicated by an arrowhead. **(E)** HECTD1 was also cleaved in the human leukaemia monocytic cell line THP-1, as soon as 30 min post STS treatment and again blocked by pre-treatment with z-VAD-FMK (Lane 7 vs. 8). **(F)** Induction of extrinsic cell death pathways following treatment of HeLa cells with TNF-α/CHX also led to a similar pattern suggestive of HECTD1 cleavage. Note that treatment with TNF-α alone, which activates NF-κB but not apoptotic cell death, had no effect on HECTD1. **(G)** HECTD1 cleavage in Hela cells upon TNF-α/CHX treatment is blocked by z-VAD-FMK (Lane 3 vs. 4). Membranes were cut to enable detection with multiple antibodies. A-C were created with https://BioRender.com.

### 4.2. HECTD1 is proteolytically processed during apoptosis

Analysis of the PROTOMAP data revealed that following a 2 h STS treatment of Jurkat T cells, two HECTD1 fragments were detected by 1D SDS PAGE, in addition to the band corresponding to full-length HECTD1. Proteomics analysis of these two cleaved fragments identified six peptides which could be mapped to the N-terminal part of the protein and up to aa1382 for band A, and four peptides from aa1961 for band B (**Figure 1C**). This suggests that upon induction of apoptosis, HECTD1 is cleaved at a single site, likely located between aa1382 and aa1961.

To validate these results, we treated HeLa cells with STS, a potent pan-kinase inhibitor of PKCα/γ/η, which is routinely used to trigger intrinsic apoptotic cell death in the absence of death receptor signalling (57). Treatment of HeLa cells with STS yielded a single new band by western blot analysis using a HECTD1 antibody raised against the first 50 residues of the protein. This cleaved form of HECTD1 (HECTD1^N^) migrated below the signal corresponding to full-length HECTD1 (HECTD1^FL^, 289 kDa) (**Figure 1D**). Based on the migration profile, we evaluated the size of this cleaved N-terminal fragment, HECTD1^N^, to be around 180 kDa. The timing of HECTD1 cleavage post treatment was very similar to that of the apoptotic marker poly(ADP-ribose) polymerase (PARP) which is an early substrate of caspase mediated cleavage during apoptosis (**Figure 1D**) (58–61). Pre-treatment of cells with z-VAD-FMK prior to STS treatment reduced the appearance of the cleaved product but increased the proportion of the full length-protein, and this was observed for both PARP and HECTD1. STS-induced HECTD1 cleavage was also observed in the monocytic cell line THP-1 isolated from the peripheral blood of an acute monocytic leukaemia patient (**Figure 1E**). Treatment of THP1 cells revealed that full length HECTD1 was completely processed 4 h post STS and furthermore, z-VAD-FMK treatment efficiently inhibited HECTD1 cleavage (**Figure 1E, lane 8 vs. 7**). HECTD1 was also prone to cleavage in the colorectal cancer cell line HCT116 and HEK293T although these cells were less sensitive to STS and higher concentrations and incubation times had to be used (**Supplementary Figure 1A & B**). However, we did not observe cleavage of HECTD1 in HeLa cells following stimulation with mitochondrial or ER stressors including FCCP, antimycin A, ionomycin, thapsigargin, or H2O2 (**Supplementary Figure 2**).

We next examined whether HECTD1 cleavage also occurred following activation of the extrinsic branches of apoptotic cell death using TNF-α/CHX co-stimulation (62). This was indeed the case, and we observed a banding pattern indicative of a single cleavage event of HECTD1 similar to the banding pattern observed following STS treatment (**Figure 1F**). Used as control, treatment of HeLa cells with TNF-α alone, which acts as inflammatory stimulus triggering pro-survival NF-κB signalling, did not induce cleavage of HECTD1 (**Figure 1F**) (63). We also observed that HECTD1 depletion had no effect on NF-κB gene expression, assayed using a NF-κB-luciferase reporter system, indicating that it does not impact on NF-κB signalling (**Supplementary Figure 3**). We recapitulated these experiments in the presence of z-VAD-FMK which effectively blocked HECTD1 cleavage (**Figure 1G**). Just like PARP, HECTD1 cleavage induced by TNF-α/CHX occurred readily, as early as 30 min post treatment. HECTD1 cleavage was also observed following combined Poly(I:C)/CHX treatment in HeLa cells which has also been shown to induce apoptotic cell death (**Supplementary Figure 1C**) (64, 65). Together, these data validate and expand on the PROTOMAP data and clearly indicate HECTD1 as *bona fide* substrate for caspase-dependent cleavage during intrinsic and extrinsic apoptotic cell death. Further, the HECTD1 proteolytic event yields an N-terminal product (HECTD1^N^) of approximately 180 kDa and a C-terminal product (HECTD1^C^) estimated at around 100 kDa. Given HECTD1 cleavage is observed following induction of apoptotic cell death by the intrinsic and extrinsic pathways, it is likely processed by an executioner caspase.

### 4.3. HECTD1 is cleaved at Asp^1664^

Similar to Itch and NEDD4, HECTD1 appears to be cleaved at a single site following activation of apoptotic cell death. We consulted the ELM and CaspDB databases to try and identify this single cleavage site within HECTD1 by predicting high-confidence caspase consensus motifs (48, 49). By filtering high-scoring potential cleavage sites against the peptide signature produced by PROTOMAP (**Figure 1C**), we were able to narrow down our search to five putative sites all located between the MIB/HERC2 domain and the BTHB domain of HECTD1 (**Figure 2A**). We then selected three candidates based on their confidence score obtained from CaspDB as well as sequence conservation. For each candidate cleavage site tested, we substituted the P1 aspartate residue with glutamate, overexpressed these constructs encoding mouse Hectd1 in Hela cells and tested the effect of STS treatment on HECTD1 cleavage (**Figure 2B-C**). HECTD1^D1664E^ was the only mutant resistant to cleavage as shown by the single band corresponding to HECTD1^FL^ in STS-treated cells. (**Figure 2C, lane 3 vs 1**). Thus, HECTD1 is proteolytically processed at the caspase consensus sequence DFLD1664↓S.

**Figure 2.**
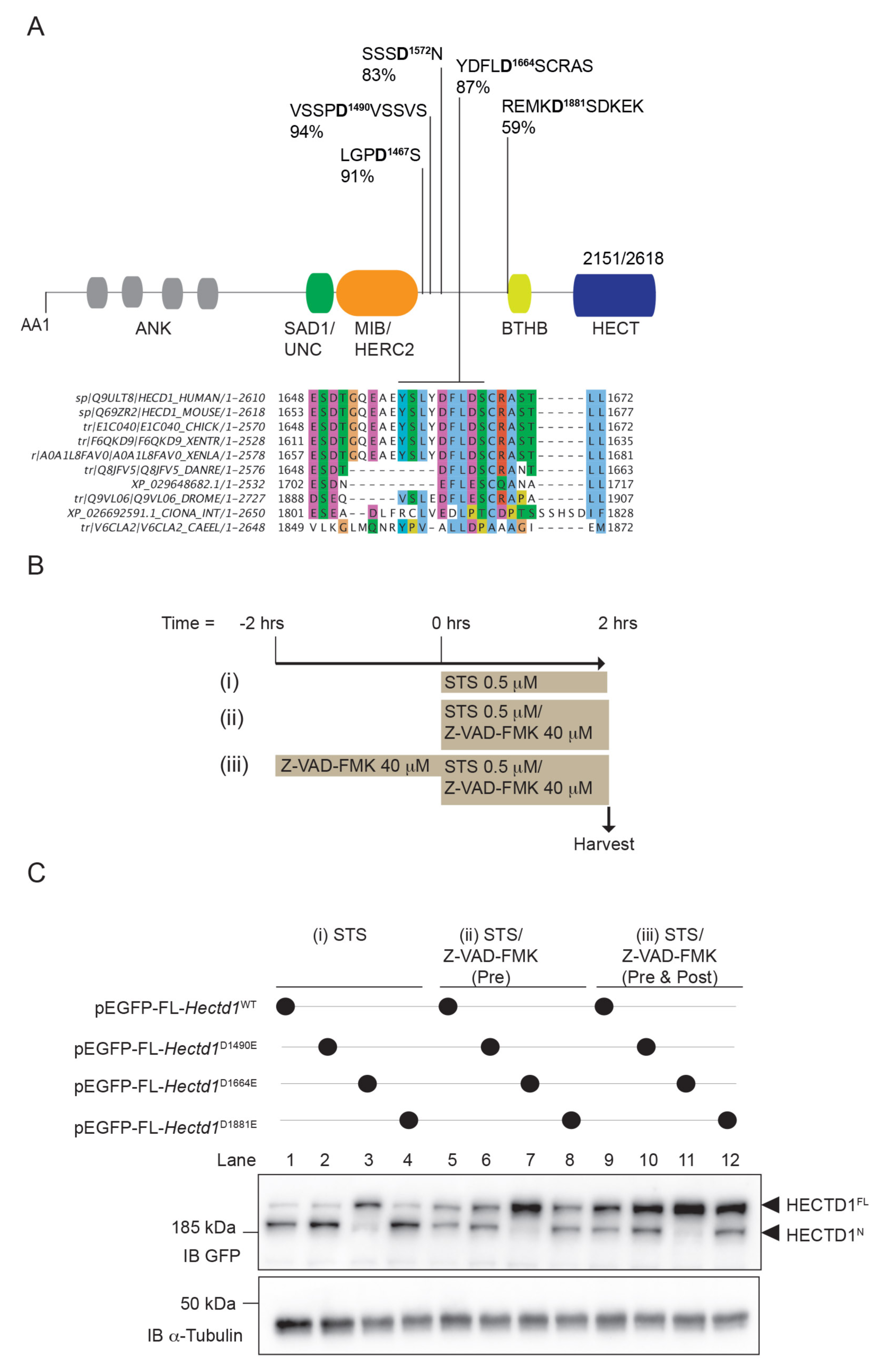
HECTD1 is cleaved at aspartate residue 1664. **(A)** Top panel: domain organisation of HECTD1 highlighting 5 putative caspase cleavage sites identified using CaspDB and located between aa1382 and aa1961 (Figure 5.1C) (Kumar et al. 2014, 2020). Lower panel: Sequence alignment showing conservation of at DFLD1664 putative caspase cleavage site. **(B)** Schematic of the treatment carried out in C. HeLa cells were treated with (i) STS only for 2 h, (ii) STS and z-VAD-FMK for 2 h, (iii) or were pre-treated with z-VAD-FMK for 2 h prior to addition of fresh inhibitor and STS for an additional 2 h. **(C)** HeLa cells were transfected with FuGENE^®^ HD for 24 h with the indicated constructs prior to treatment as shown in B. pEGFP plasmids (Clontech Laboratories, Mountain View, CA, USA) encoding full-length mouse Hectd1 with a single D to E point mutation at either D1490, D1664 or D1881 were tested. All conditions were harvested and lysed at the same time. Samples were lysed in RIPA run on 4-12% Bis-Tris PAGE and probed for HECTD1 (Ab101992) and the loading control GAPDH. Membranes were cut to enable detection with multiple antibodies.

### 4.4. HECTD1 is cleaved by caspase-3

The DFLD cleavage motif we identified in HECTD1 is reminiscent of that recognised by effector caspases 3/7, DXXD↓A/G/N/S/T, further suggesting caspases 3/7 cleave HECTD1 (66, 67). To validate this experimentally, we transiently expressed 3xFLAG-Full-length mouse HECTD1 in HEK293T cells, followed by FLAG IP using anti-FLAG M2 magnetic beads and incubation of the washed beads with 1 unit of each of the 10 recombinant caspases (**Figure 3A**). Although we observed cleaved fragments when using all caspases, only incubation with recombinant caspase-3 produced a convincing pattern reminiscent of the single cleavage event we observed in **Figure 1**. To further validate this, we also tested 3xFLAG-FLmHECTD1^D1664E^ and as expected we observed this construct could not be cleaved by recombinant caspase-3, as shown by the absence of the C-ter HECTD1 fragment observed in Figure 3A (**Figure 3B, Lane 4**). Cleaved products obtained with incubation with other caspases indicate that in the presence of excess recombinant active caspases, HECTD1 can be cleaved at sites other than DFLD1664↓S. This is not surprising and most likely reflect the limitation of such *in vitro* assay.

**Figure 3.**
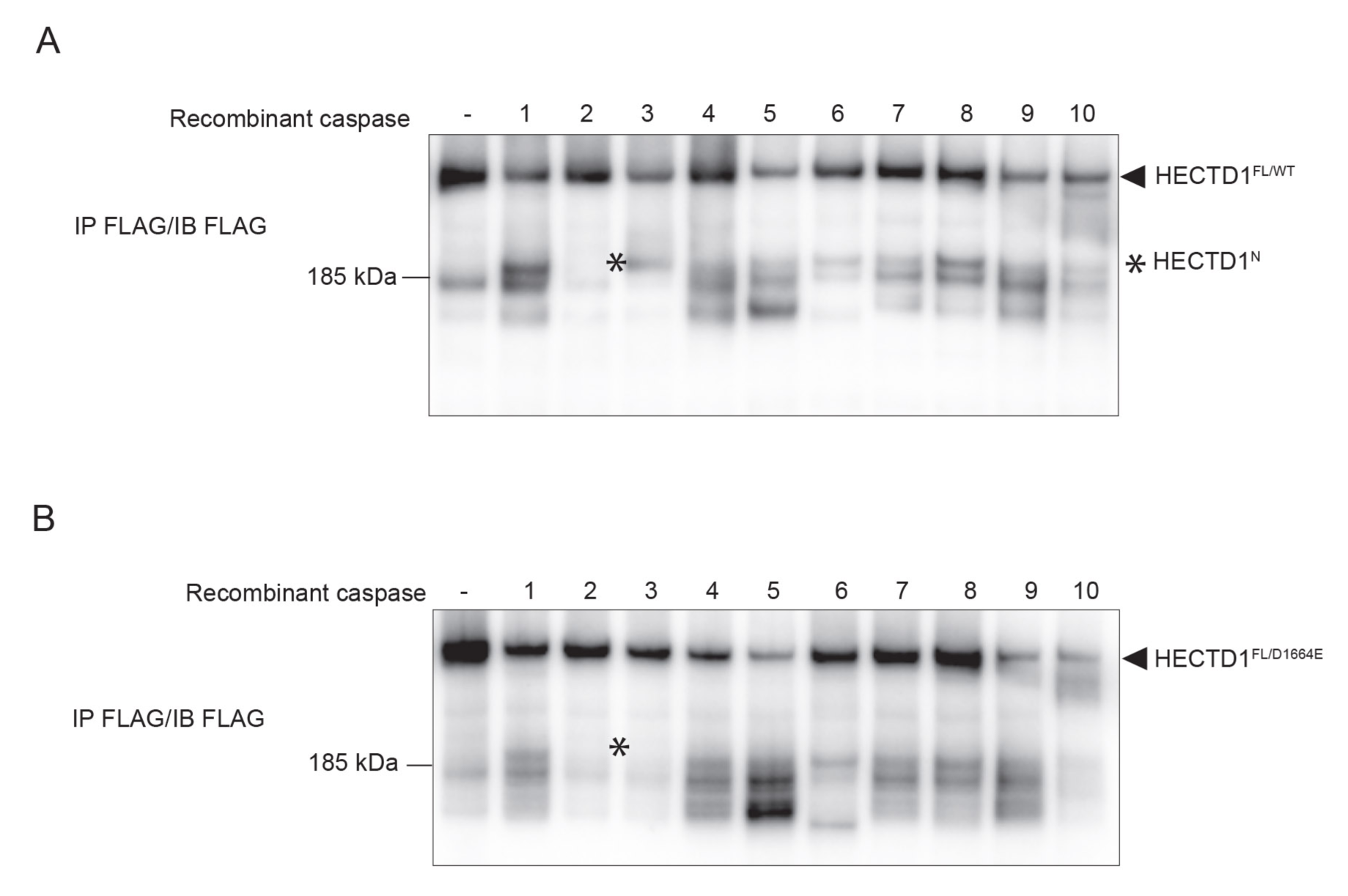
HECTD1 is cleaved by caspase-3. **(A)** *In vitro* caspase cleavage assay using 3xFLAG-FL-mouseHectd1. To determine which caspase was responsible for HECTD1 cleavage, HEK293T cells were transfected with 3xFLAG-FLmHectd1 using Lipofectamine 2000 for 24 h followed by immunoprecipitation with Dynabeads magnetic beads. Beads were resuspended in buffer and aliquoted. Each aliquot of 3xFLAG-FLmHECTD1WT was incubated with 1 unit of each recombinant caspases as indicated, for 1 h at 37°C followed by addition of 2X LDS/100 mM DTT to terminate reactions. Samples were resolved on 3-8% Tris-Acetate gels and probed with anti-Flag antibody. **(B)** Same as in A but with 3xFLAG-FLmHECTD1^D166E^, in order to highlight cleavage events not targeting the DFLD1664 motif. Asterisk (*) indicates the cleaved HECTD1 bands produced upon induction of apoptotic cell death which is not produced when using the D1664E mutant. Membranes were cut to enable detection with multiple antibodies.

To further confirm HECTD1 cleavage is specifically induced by caspase-3 during apoptotic cell death and not via other pathways driven by the activation of other caspases, we investigated the effect of pyroptosis on HECTD1 cleavage. Pyroptosis is triggered through sensing of pathogen-and danger-associated molecular patterns (PAMPs and DAMPs) including viral dsRNA, by pattern recognition receptors (PRRs) such as the Toll-like receptor TLR3 (68, 69). Pyroptosis triggered by dsRNA sensing drives the assembly of the NLRP3-inflammasome which enables the maturation of IL-18 and IL1-β through caspase 1-mediated proteolytic processing. Alternatively, a non-canonical inflammasome pathway can be induced by LPS. Here, intracellular LPS is sensed directly by caspase 4/5/11 which in turn become activated, cleave the pore-forming effector protein Gasdermin D (GSDMD), and mediate secondary activation of the NLRP3 inflammasome and caspase 1 (70). To further exclude the possibility that HECTD1 may also be proteolytically processed by these caspases, we tested the effect of non-transfected or transfected LPS with the latter being a potent activator of pyroptosis. However, we found that in THP-1 or PMA-differentiated THP-1 cells which acquire a macrophage-like phenotype, HECTD1 was not cleaved (**Supplementary Figure 4)**. Together these data indicate that caspase-3 is the sole caspase responsible for HECTD1 cleavage at DFLD↓S during apoptotic cell death.

### 4.5. The HECTD1^C^ cleaved product is stable

Cleavage of Itch at Asp^240^ releases its N-terminal C2 domain, and ΔC2-Itch retains ubiquitin ligase activity towards its canonical substrate ΔNp63α (24). In contrast, caspase-1 mediated cleavage of the RBR E3 ubiquitin ligase PARKIN inhibits its ligase activity and contributes to inflammasome-mediated block of mitophagy by removing the N-terminal ubiquitin-like domain (22,29,71). In the case of HECTD1, we also observe cleavage in an inter-domain region which similarly to Itch separates N-terminal domains from the C-terminal catalytic HECT domain. Since the HECT domain remains intact and thus retains the potential for catalytic activity, we set out to compare the stability of the C-terminal cleavage fragment of HECTD1 (HECTD1^1665-2610^) to that of endogenous full-length HECTD1 as an initial indicator of functional potential (**Figure 4**).

**Figure 4.**
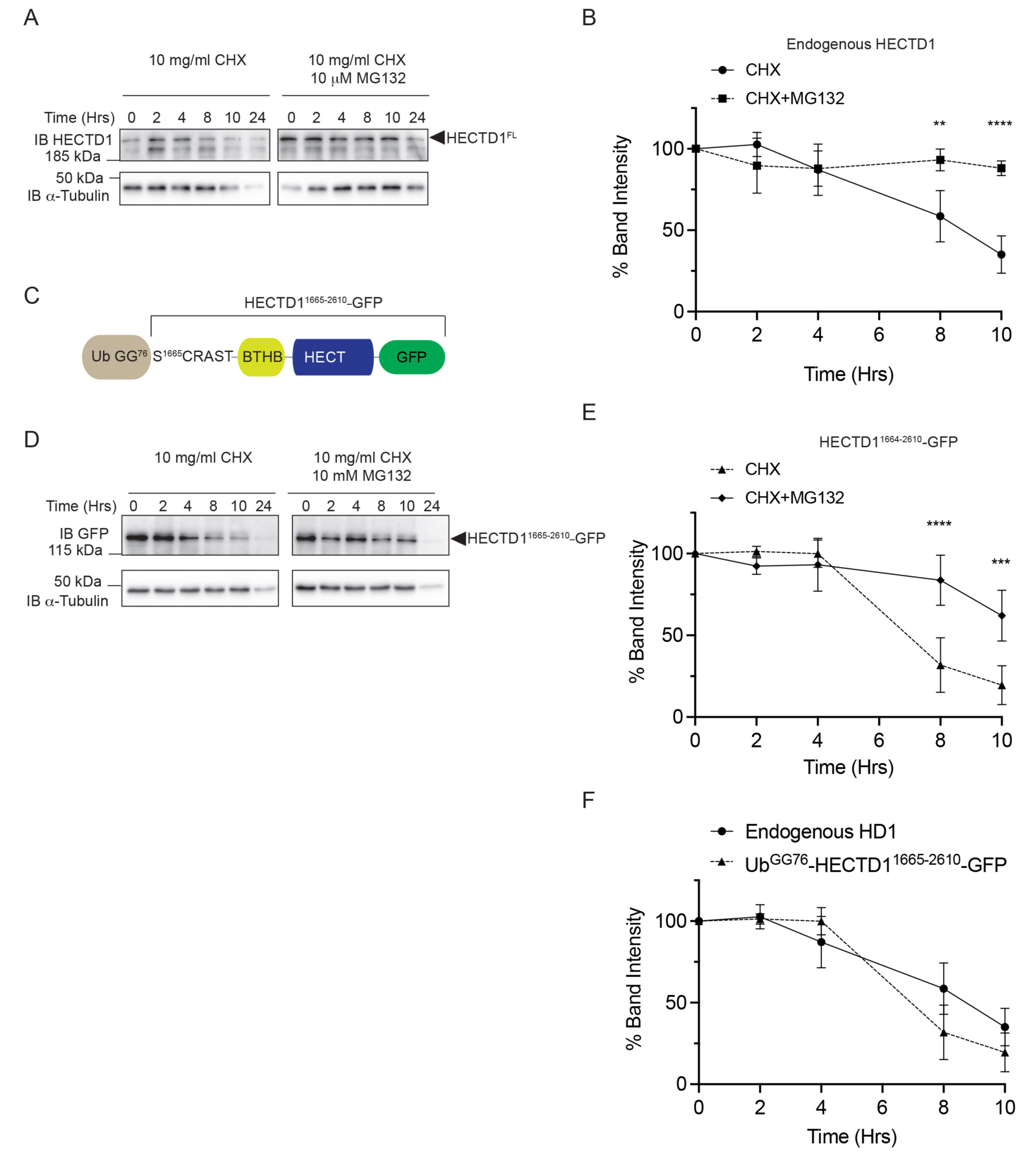
HECTD1C-terminal fragment (aa1665-2610) has a similar stability to full-length endogenous HECTD1. **(A)** CHX chase experiments were carried out in HEK293T cells to determine the stability of endogenous HECTD1. HEK293T cells were treated with 10 µg/ml CHX either alone or in combination with 10 µM MG132. Samples were harvested at the indicated time points, lysed in RIPA and analysed by immunoblotting using HECTD1 antibody and GAPDH antibody which was used as loading control. Data shown is representative of at least three experiments. **(B)** Quantification of gel images shown in A (n = 3 independent experiments). Signal intensity was measured using ImageJ, the ratio for HECTD1/Tubulin quantified, and then analysed for statistical significance using 2-way ANOVA with Šídák’s multiple comparisons test; ****p<0.0001, ***p<0.001, **p<0.01, and *p<0.05. **(C)** Cartoon showing the plasmid construction to determine the stability of Hectd1^1665-2610^. The design is inspired by work from Varshavsky and colleagues who used it to determine the stability of N-terminal residues to define the N-end rule pathway (Passmore et al. 1986; Bachmair and Varshavsky 1989). Briefly, ubiquitin was cloned upstream of P1’ Ser^1665^. Upon translation, ubiquitin is co-translationally cleaved by endogenous DUBs thereby revealing the N-terminal residue in this case Ser^1665^. A GFP tag located at the C-terminus enables the detection of this HECTD1^1665-2610^ protein. **(D)** UbGG^76^-HECTD1^1665-2610^-GFP was transiently transfected in HEK293T cells and CHX chase was carried out as in A. **(E)** Data shown were analysed as in A using ImageJ and %band intensity is shown relative to the corresponding alpha-Tubulin signal. MG132 was used to show that the decrease in protein levels is due to protein turnover by the 26S proteasome. Membranes were cut to enable detection with multiple antibodies.

We first used a cycloheximide (CHX)-chase assay to ascertain the stability of HECTD1 which had a half-life of around 8 h and as a control we also showed that HECTD1 levels could be stabilised in the presence the proteasomal inhibitor MG132 (**Figure 4A & B**). We next determined the stability for the C-terminal cleaved product of HECTD1 (HECTD1^C^) A key determinant of protein fate following protease processing is the Arg/N-end rule pathway which established serine is not a destabilising N-degron, and therefore the newly exposed HECTD1 P1’ residue Ser^1665^ likely suggests this cleaved fragment will be stable (72). In order to examine the stability of HECTD1^C^, we utilised the ubiquitin-fusion strategy whereby a ubiquitin moiety is fused N-terminally of the residue of which stability is being tested, here Ser^1665^. Expression of this construct in mammalian cells triggers co-translational deubiquitylation of the protein and thus removal of the N-terminally fused ubiquitin moiety at Gly^76^ by endogenous DUBs, exposing the N-terminal Ser^1665^ residue (**Figure 4C-E**) (72). The CHX chase assay confirmed mouse HECTD1^C^ exhibits a comparable half-life to human endogenous full-length HECTD1 (**Figure 4F**).

### 4.6. HECTD1 positively contributes to extrinsic and intrinsic apoptotic cell death

We next pondered what could be the role, if any, HECTD1 cleavage during apoptotic cell death by caspase-3. Therefore, we set out to further examine the significance of these data by exploring the effect of HECTD1 depletion on apoptotic cell death (**Figure 5**). Transient HECTD1 depletion in Hela cells treated with TNF-α/CHX (extrinsic apoptotic pathway) or STS (intrinsic apoptotic pathway) resulted in a marked decrease in the apoptotic cell population (Annexin V^+^ DAPI^-^) in both, the HECTD1 SMARTPool and HECTD1 siRNA#6-treated conditions (**Figure 5A-C**). Following TNF-α/CHX treatment, on average 18% (HECTD1 SMARTPool) and 26% (HECTD1 siRNA #6) less apoptotic cells were observed compared to a non-targeting (NT) siRNA control, whilst a more dramatic decrease of 37% (HECTD1 SMARTPool) and 46% (HECTD1 siRNA #6) was observed following STS treatment (**Figure 5C**). This suggests that HECTD1 plays a pro-apoptotic role in both the intrinsic as well as extrinsic apoptotic pathways. To explore how HECTD1 depletion might affect cell death, we employed a multiplex fluorescent assay to monitor caspase-3, 8 and 9 activity in HECTD1 depleted cells upon TNF-α/CHX (**Figure 5D & E**) or STS (**Figure 5F & G**) treatment in HeLa cells. During both intrinsic or extrinsic apoptotic cell death, HECTD1 depletion significantly reduced Caspase-3, but not caspase 8 or 9 activity, compared to the controls. Our findings are in line with data indicating that HECTD1 stable depletion in T47D breast cancer cell lines does not affect caspase-3 protein levels but rather its cleavage (33).

**Figure 5.**
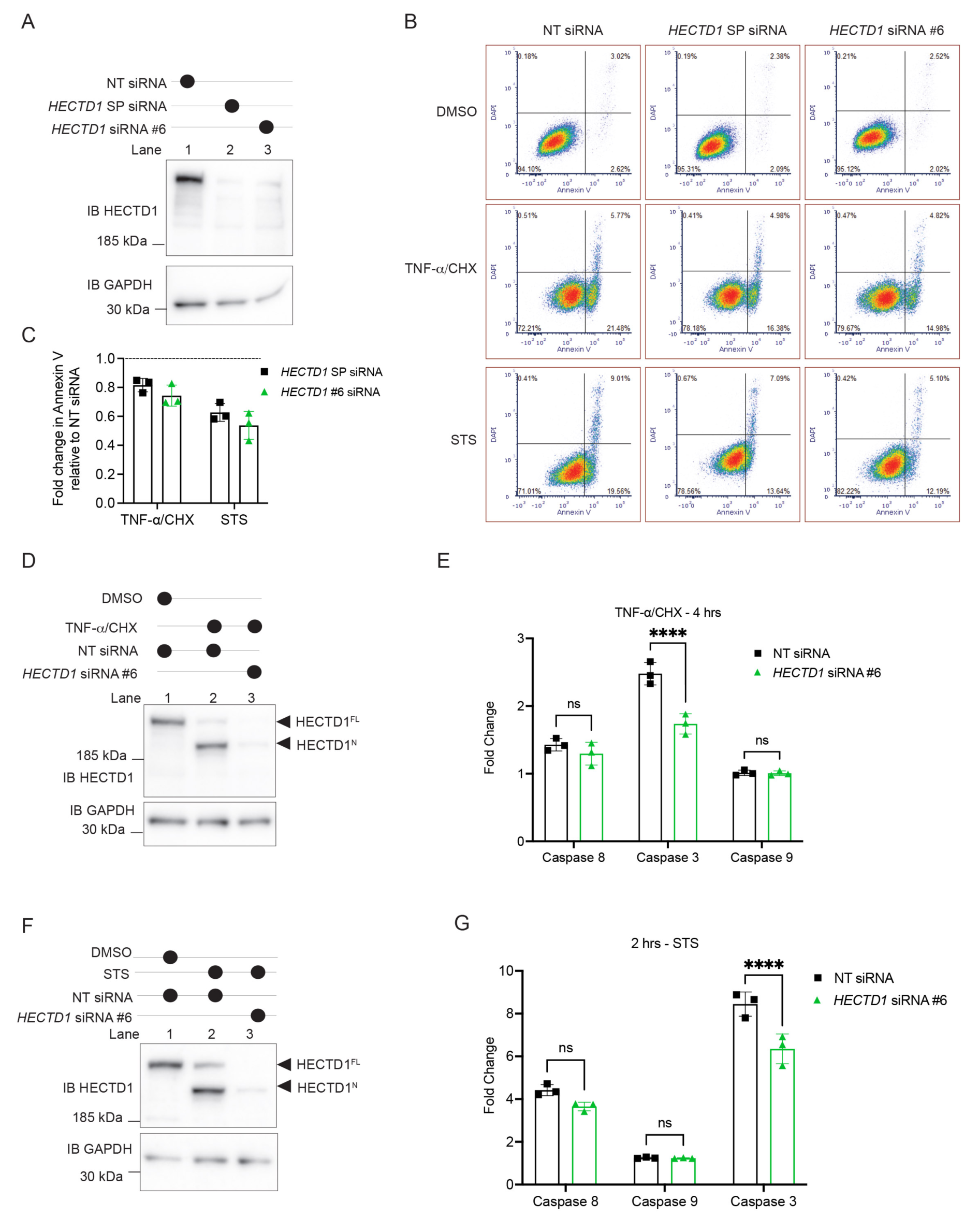
HECTD1 is a positive component of apoptotic cell death. **(A)** Immunoblot showing the extent of HECTD1 depletion in Hela cells prior to induction of apoptosis and measurement of apoptotic cell death by flow cytometry. **(B)** Measuring apoptotic cell population using flow cytometry quantification of Annexin V^+^/DAPI^-^ staining. HeLa cells were transfected using RNAiMAX with either non-targeting siRNA, HECTD1 SMARTPool siRNA or individual HECTD1 siRNA #6 for 48 h prior to treatment with DMSO, STS (0.5 µM) for 2 h, or TNF-α (50 ng/ml) and CHX (10 µg/ml) for 4 h. Following treatment, cells were harvested and resuspended in 1X Annexin V binding buffer containing Annexin V-Alexa Fluor 555 conjugate and 300 nM DAPI. Following a 15 min incubation, samples were placed on ice and analysed using a BD FACSAria III immediately. Data were analysed using FCS Express 7 Research Edition. Flow cytometry data shown is representative of n = 3 independent experiments. **(C)** Fold change in apoptotic cell population compared to NT-siRNA condition is shown for STS and TNF-α + CHX treatments. **(D)** Immunoblot showing the extent of HECTD1 depletion in Hela cells following induction of apoptosis with TNF-α (50 ng/ml) and CHX (10 µg/ml) and prior to measurement of caspase-3, caspase-8 and caspase-9 activity using the Multiplex Activity Assay Kit. **(E)** Hela cells were transfected with either non-targeting siRNA, HECTD1 SMARTPool siRNA or individual HECTD1 siRNA #6 for 48 h prior to treatment with either DMSO, or TNF-α (50 ng/ml) and CHX (10 µg/ml) for 4 h. Following treatment, cells were harvested, lysed in RIPA, and samples were then analysed by immunoblotting using HECTD1 antibody and GAPDH which was used as loading control. Quantification of fluorescence for caspase-3, caspase-8 and caspase-9 was carried out according to the manufacturer’s protocol using a GloMax®-Multi Detection System. **(F)** Immunoblot showing the extent of HECTD1 depletion in Hela cells following induction of apoptosis with STS (0.5 µM) for 2 h and prior to measurement of caspase-3, caspase-8 and caspase-9 activity using the Multiplex Activity Assay Kit. **(G)** Transfections and fluorescent measurement of caspase-3, caspase-8 and caspase-9 were carried out and measured as in E. For each experiment, data were measured in triplicate and data are representative of n = 2 independent experiments. Fold change is relative to the DMSO condition which was set as 1. Statistical analysis was carried out using 2-way ANOVA with Šídák’s multiple comparisons test; ****p<0.0001, ***p<0.001, **p<0.01, and *p<0.05. Membranes were cut to enable detection with multiple antibodies.

To try and address the functionality of HECTD1 cleavage during apoptotic cell death, we tested whether the caspase resistant construct mouse Hectd1^D1664E^ might rescue the decrease in cell death observed upon HECTD1 depletion (**Figure 6**). HeLa cells were treated for 48 h with NT or HECTD1 siRNA#8 prior to STS treatment to deplete endogenous HECTD1 levels. We used HECTD1 siRNA#8, since full-length mouse Hectd1 constructs are refractory to the previously used HECTD1 siRNA#6 (**Figure 6A & B**). We found that re-expression of the caspase resistant Hectd1 construct (3xFLAG-FLmHectd1^D1664E^) but not the double mutant (3xFLAG-FLmHectd11^D1664E/C2579G^) which is resistant to caspase cleavage and catalytically inactive, rescued the decrease in apoptotic cell population (Annexin V^+^ DAPI^-^) observed upon HECTD1 knockdown (**Figure 6C**). We also tested this in our caspase multiplex fluorescent assay system and found that neither constructs had an effect on caspase 8 nor 9 activity (**Figure 7A-D**). In contrast, HECTD1^D1664E^ but not HECTD1^D1664E/C2579G^ rescued the decrease in caspase-3 activity triggered by HECTD1 transient depletion (**Figure 7E**). Collectively, these data suggest full-length HECTD1 is required to contribute its positive effect on apoptotic cell death, likely through its ubiquitin ligase activity, and does so via a positive effect which results in caspase-3 activation.

**Figure 6.**
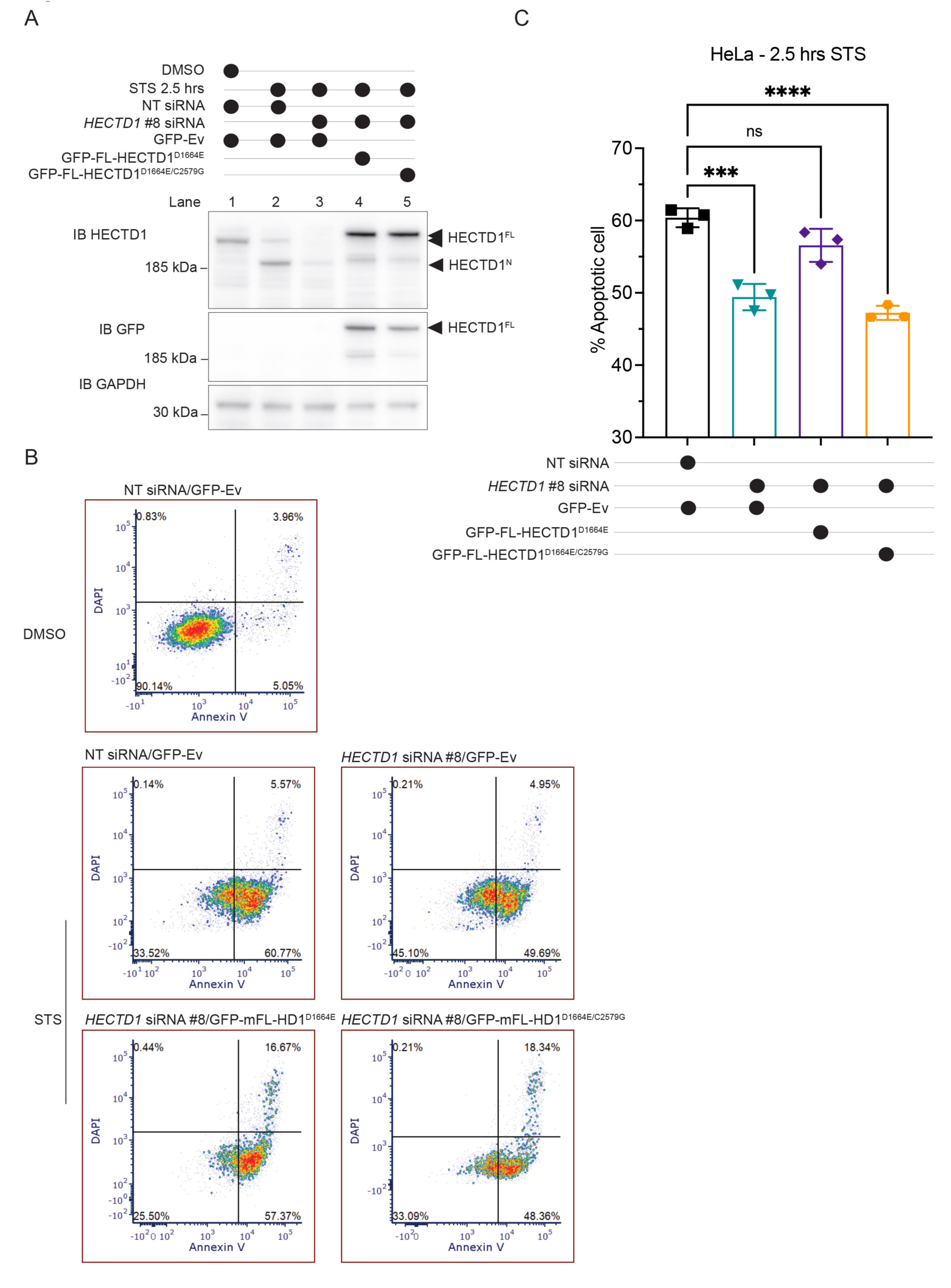
Full-length HECTD1 is required for its effect on caspase-3 activity. **(A)** Immunoblot analysis showing the efficiency of HECTD1 knockdown (Lane 3), and the re-expression of cleavage resistant constructs (Lane 4 and 5). HeLa cells were transfected using RNAiMAX with either non-targeting siRNA, HECTD1 SMARTPool siRNA or individual HECTD1 siRNA #8 for 48 h prior to treatment; and with either GFP-empty vector, GFP-full-length mouse HECTD1^D1664E^, or GFP-full-length mouse HECTD1^D1664E/C2579G^ 24h prior to treatment with DMSO, STS (0.5 µM) for 2 h. **(B)** Assessment of cell death using flow cytometry quantification of Annexin V/DAPI staining. Cells were transfected and treated as in A. Following treatment, cells were harvested and resuspended in 1X Annexin V binding buffer containing Annexin V-Alexa Fluor 555 conjugate and 300 nM DAPI. Following a 15 min incubation, samples were placed on ice and analysed using a BD FACSAria III immediately. Data were analysed using FCS Express 7 Research Edition. **(C)** Quantification of data shown in B. Data are representative of n = 3 independent experiments. Statistical analysis was carried out using one-way ANOVA with Dunnett’s multiple comparisons test; ****p<0.0001, ***p<0.001, **p<0.01, and *p<0.05. Membranes were cut to enable detection with multiple antibodies.

**Figure 7.**
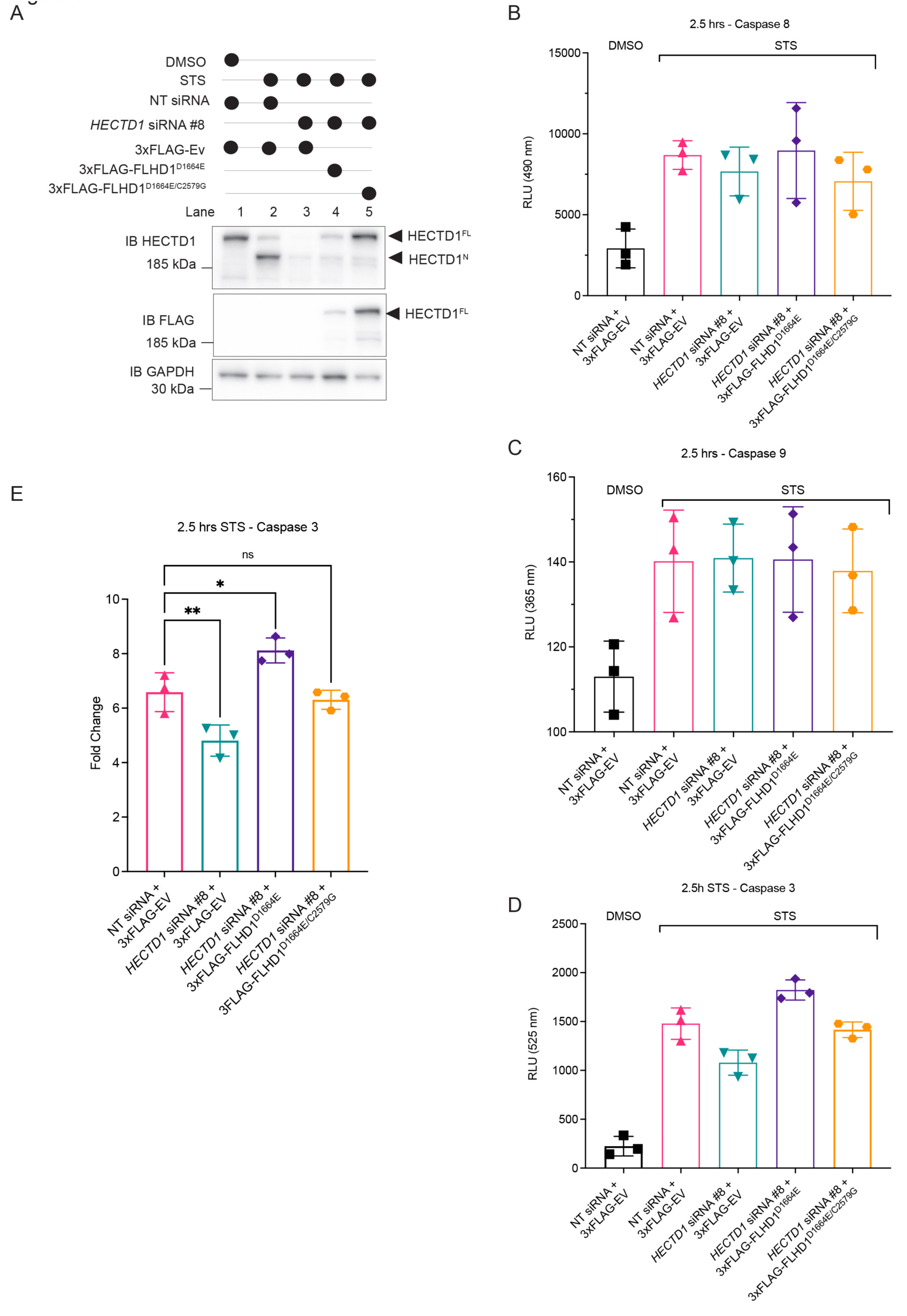
Full length HECTD1 contributes to caspase-3 proteolytic activity through its catalytic activity. **(A)** Immunoblot analysis showing the efficiency of HECTD1 knockdown (Lane 3), and the re-expression of cleavage resistant constructs (Lane 4 and 5). Note that 3xFLAG-Full-length mHectd^D1664E/C2579G^ is more stable than 3xFLAG-Full-length mHectd1^D1664E^ (Lane 5 vs. 4). HeLa cells were transfected using RNAiMAX with either non-targeting siRNA, HECTD1 SMARTPool siRNA or individual HECTD1 siRNA #8 for 48 h prior to treatment; and with either 3xFLAG-empty vector, 3xFLAG-full-length mouse Hectd1^D1664E^, or 3xFLAG-full-length mouse HECTD1^D1664E/C2579G^ 24h prior to treatment with DMSO, STS (0.5 µM) for 2 h. **(B-D)** Cells were transfected and treated as in A. Quantification of fluorescence for caspase-3, caspase-8 and caspase-9 (Multiplex Activity Assay Kit) was carried out according to the manufacturer’s protocol using a GloMax®-Multi Detection System. **(B)** Relative light units for 490 nm which measure caspase 8 activity. **(C)** Relative light units for 365 nm which measure caspase 9 activity. **(D)** Relative light units for 525 nm which measure caspase-3 activity. **(E)** Fold change for caspase-3 activity. For each experiment, data were measured in triplicate and data are representative of n = 2 independent experiments. Fold change is relative to the DMSO condition which was set as 1. Statistical analysis was carried out using one-way ANOVA with Dunnett’s multiple comparisons test; ****p<0.0001, ***p<0.001, **p<0.01, and *p<0.05. Membranes were cut to enable detection with multiple antibodies.

### 4.7. TRABID UBD identifies atypical ubiquitin chains during apoptotic cell death

To further explore HECTD1’s putative new role in apoptosis, we exploited the ubiquitin binding domain of the deubiquitinating enzyme TRABID. We recently reported TRABID-HECTD1 as DUB-E3 pair handling ubiquitin chains containing K29 linkages (45). The TRABID NZF was shown to preferentially bind to and cleave K29 and K33 linkages over other linkages including K63 (44,46,73,74). Further, we recently reported that the HECTD1 HECT domain assembles K29- or K48-linked chains and that full E3 ligase activity involved branched K29/K48 ubiquitin linkages (45). Importantly our data also showed that the TRABID NZF-1-3 domains efficiently pulled down the chains assembled by HECTD1. Therefore, we next used TRABID as a tool to further illuminate the “ubiquitin code” during cell death (**Figure 7**).

We used GST-TRABID NZF 1-3 (aa1-200) as bait on apoptotic cell lysates and found that TRABID NZF 1-3 baits trapped more ubiquitin species from HeLa cells treated with TNF/CHX and to a lesser extent STS, than DMSO treated cells (**Figure 7A, lane 9 and 8 vs 7**). These ubiquitin signals are likely composed of K29 and K33 ubiquitin linkages given TRABID’s preference for these linkages. However, and since TRABID GST-TRABID NZF 1-3 has some ability to bind K63-linked chains, we also employed the K63-specific NZF domain of TAB2 as control (44, 75). However, we found that this GST-TAB2 bait was less efficient at enriching for ubiquitin species in apoptotic cells (**Figure 7A, lane 6 and 5 vs 4**). A TRABID NZF 1-3 ubiquitin binding deficient mutant, which has the three NZF domains mutated at TY>>LV, was used alongside GST as control and no ubiquitin signal was detected (**Figure 7A**). We can in part attribute this enrichment in ubiquitin signals following induction of apoptosis to HECTD1 ubiquitin ligase activity since this signal was markedly reduced upon HECTD1 transient knockdown (**Figure 7B, lane 12 vs. 10**). In contrast, there was no change in the signal obtained using the K63-binder GST-TAB2 (**Figure 7B, lane 8 vs 6**). This data suggests ubiquitin chains, recognised by TRABID and contributed by HECTD1 ligase activity and thus likely containing K29 linkages, are increased during apoptotic cell death.

## 5. Discussion

The regulation of protein fate and function is well understood from a transcriptional and post-translational regulation standpoint. Although protein cleavage is usually associated with the digestion of proteins in lysosomes, or by the proteasome, such events can be part of functional mechanisms and contribute to the regulation of cellular processes. The caspase family is a prime example, with caspases synthesised as inactive zymogens which need to be cleaved in order to gain their proteolytic activity towards their substrates. In contrast to this activating mechanism, the proteolytic cleavage of some E3 ubiquitin ligases has been linked to loss of ubiquitin ligase activity, including for cIAP1/2, XIAP, MDM2, RBX1, HOIP and HOIL1 (18,21,22,29,76–78). Cleavage of HECT E3 ubiquitin ligases on the other hand does not seem to affect protein ubiquitylation per se and instead might affect substrate recruitment (24). Interestingly, these cleavage events occur at a single site and in flexible interdomain regions releasing intact and potentially functional domains.

HECT E3 ubiquitin ligases have been proposed to exist in an inactive conformation which would indicate these enzymes can readily deploy their ubiquitin ligase activity in response to the appropriate cellular cues. This is the case for Itch and NEDD4 for with the N-terminal C2 domain interacts with the catalytic HECT domain in such a way that the active site is occluded or by blocking E2-E3 trans-thiolation. In addition to the C2 domain, WW domains and linker regions located N-terminally from the HECT domain have also been shown to contribute to autoinhibition of ubiquitin ligase activity for NEDD4 and Itch (79). Deletion of the WW2L-HECT interface activates Itch activity *in vitro*, further arguing that NEDD4-type E3s are in fact pro-E3 ubiquitin ligases. Both Itch and Nedd4 were found to be cleaved at single aspartate residues releasing the N-terminal C2 domain from the rest of the protein, composed of WW domains and the catalytic HECT domain (19, 24). *In vitro* assays have revealed that both full-length and ΔC2 Itch can ubiquitinate its canonical substrate ΔNp63α, although these assays were not quantitative. These data suggest that cleaved Itch is *in vitro* at least, an active E3 ubiquitin ligase and raises the question of whether ΔC2 Itch might have “neo” function and substrates *in vivo*.

In this study we investigated whether additional E3 ubiquitin ligase might be prone to proteolytic cleavage. We searched the PROTOMAP database for E3 ubiquitin ligases with evidence of a single cleavage site which would not affect the integrity of the catalytic HECT domain, since our thinking was that this could serve as proxy for potential functionality. PROTOMAP analysis validated NEDD4 and Itch and further revealed HECTD1 as third HECT E3 ligase fulfilling our criteria (Dix et al. 2008). We validated that HECTD1 is cleaved upon intrinsic or extrinsic apoptotic cell death in multiple cells lines including HeLa, THP-1 human monocytes, THP-1 cells differentiated into macrophage-like cells, HCT116 and HEK293T cells. We identified DFLD1664↓S as the single cleavage motif within HECTD1 which is targeted by caspase-3 activity. HECTD1’s DFLD motif and the P1’ residue (Ser^1664^ in human) are fully conserved in most vertebrate chordates we examined but not in invertebrate chordates including *Ciona intestinalis* (EDLP), *Caenorhabditis elegans* (ALLD), *Drosophila melanogaster* (DFLE) or in *Octopus vulgaris* (EFLE).

Given that extrinsic and intrinsic apoptotic cell death converge at caspase-3 activation and thus HECTD1 cleavage, we explored whether HECTD1 might play a functional role during cell death. We indeed found that HECTD1 is a positive regulator of apoptotic cell death and our data revealed that this is likely mediated through an effect on caspase-3 protease activation and/or activity. Although our data did not yet establish whether HECTD1 directly regulates caspase-3, there are precedents for feedback loops between ubiquitin ligase function and caspase activity in Drosophila and Arabidopsis (80, 81). Although there is no such link between caspase-dependent cleavage of E3 ubiquitin ligases and caspase activation/activity in human, a recent study showed that the inactivation of TRAIL-induced caspase 8 by K63-linked polyubiquitylation of caspase 8 Lys^215^ is contributed by HECTD3 ubiquitin ligase activity (82).

Although linear K63 and branched K48/K63 chains have been shown to contribute to NF-κB signalling, the ubiquitin landscape during apoptotic cell death is less well understood (83). To address this, we used ubiquitin binding domains of TAB2, a K63-specific UBD, and TRABID NZF1-3 which preferentially traps K29- and K33-linked ubiquitin chains over any other chain type including K63 chains (73, 74). We detected an enrichment of ubiquitin chains trapped by TRABID NZF 1-3 upon apoptotic cell death and established that some of this increased signal was contributed by HECTD1 ubiquitin ligase activity. We therefore propose a model whereby HECTD1 actively contributes to apoptotic cell death, likely through an increase in K29- containing linkages as this is one of the favoured ubiquitin linkages recognises by TRABID NZF. In turn, this leads to caspase-3 activation and cleavage of HECTD1 at Asp^1664^ which could serve as feedback mechanism (**Figure 9**). The E3 ubiquitin ligase cIAP1 modifies caspase-3 with degradative K48-linked polyubiquitin chains, and it will be important to establish whether HECTD1 also directly regulates caspase-3 activity through its ubiquitin ligase activity as well as the type of ubiquitin linkages involved (84). We recently established that *in vitro* at least, the isolated HECT domain of HECTD1 preferentially assembles K29- and K48-containing linkages and these can exist in a branched topology (45). It would also be interesting to compare the processivity and chain specificity of HECTD1 ubiquitin ligase activity in the context of full-length HECTD1 versus the cleaved C-ter HECTD1 fragment produced through caspase-3 cleavage (HECTD1^C^). Our findings that HECTD1 is proteolytically processed by caspase-3 is particularly interesting given Duhamel and colleagues reported a reduction in apoptotic gene expression via gene set enrichment analysis in HECTD1-depleted T47D breast cancer cells treated with the cisplatin (33). Importantly, the authors also showed that T47D-HECTD1 shRNA stable cells showed impaired activation of caspase-3. This set of data, together with our findings, might indeed suggest HECTD1 actively contributes to apoptotic cells death through the regulation of caspase-3 activity but perhaps too through the transcriptional regulation of anti-and pro-apoptotic gene expression. It will be interesting to further define molecular mechanisms regulated by HECTD1 and which may contribute to its function during cell death in order to refine our model.

**Figure 8.**
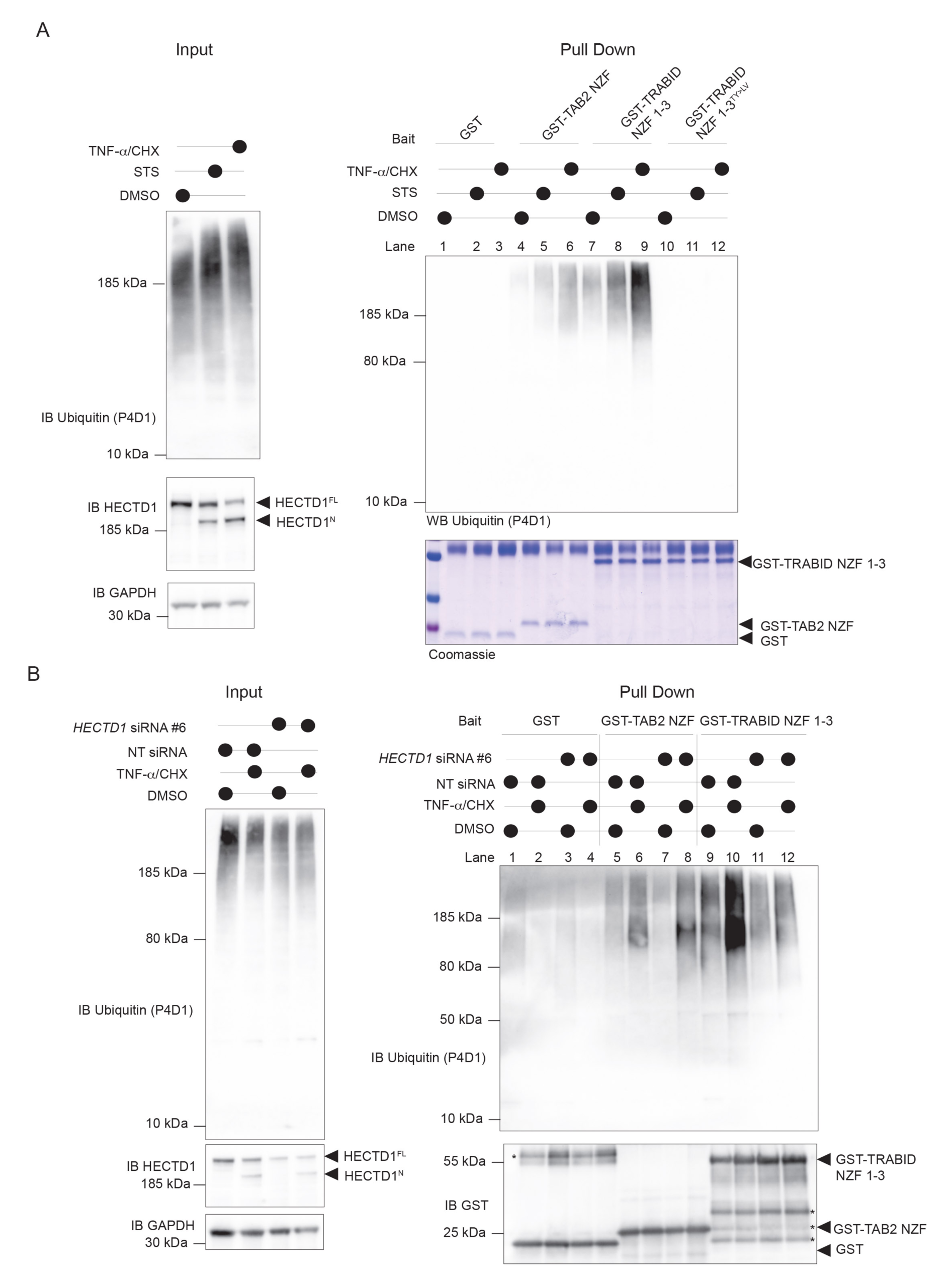
Enrichment of non-canonical ubiquitin chains during apoptosis. **(A)** TRABID NZF 1-3 preferentially bind ubiquitin chains assembled via K29 and K33 linkages over any other ubiquitin linkages (Kristariyanto et al. 2015; Michel et al. 2015). Since TRABID can also bind K63 linkages we used TAB2 NZF as control which exclusively binds K63 linkages (Komander et al. 2009; Kanayama et al. 2004). HEK293T cells were treated with either DMSO, STS for 2 h (0.5 µM), or a combination of TNF-α (50 ng/ml) and CHX (10 µg/ml) for 4 h. Following treatment, cells were pelleted and lysed in RIPA buffer supplemented with protease inhibitor, 10 mM iodoacetamide and 2 mM NEM. Inputs are shown on the left-hand side and represent 2% of the overall sample which was used for the pulldown. Samples were probed for ubiquitin using the P4D1 clone, HECTD1 (Ab101992) and GAPDH as loading control. Right. GST pulldowns were carried out as previously described (Harris et al. 2021). To capture ubiquitin species, we used 10 µg of the indicated GST-tagged protein per condition, including GST, GST-TAB2 NZF (K63 binder), GST-TRAB NZF 1-3 (K29/K33>>>K63). We also included a ubiquitin binding-deficient mutant where key TY residues in each of the three NZF have been mutated to LV (Tran et al. 2008). Coomassie stained gels show the loading of GST baits for each condition. **(B)** The increase in ubiquitin signals captured by GST-TRABID NZF 1-3 is contributed, at least in part, by HECTD1. HeLa cells were transiently transfected with either a non-targeting (NT) siRNA or HECTD1 #6 siRNA using RNAiMAX for 48 h prior to treatment with TNF-α (50 ng/ml) and CHX (10 µg/ml) for 4 h, and DMSO was used as control. Left, shows input for ubiquitin and HECTD1 (2%). Right, pulldown was carried out as in A. The membrane was also probed with a GST-tagged antibody. Asterisk (*) indicate degradation products corresponding to individual NZF. Membranes were cut to enable detection with multiple antibodies.

**Figure 9.**
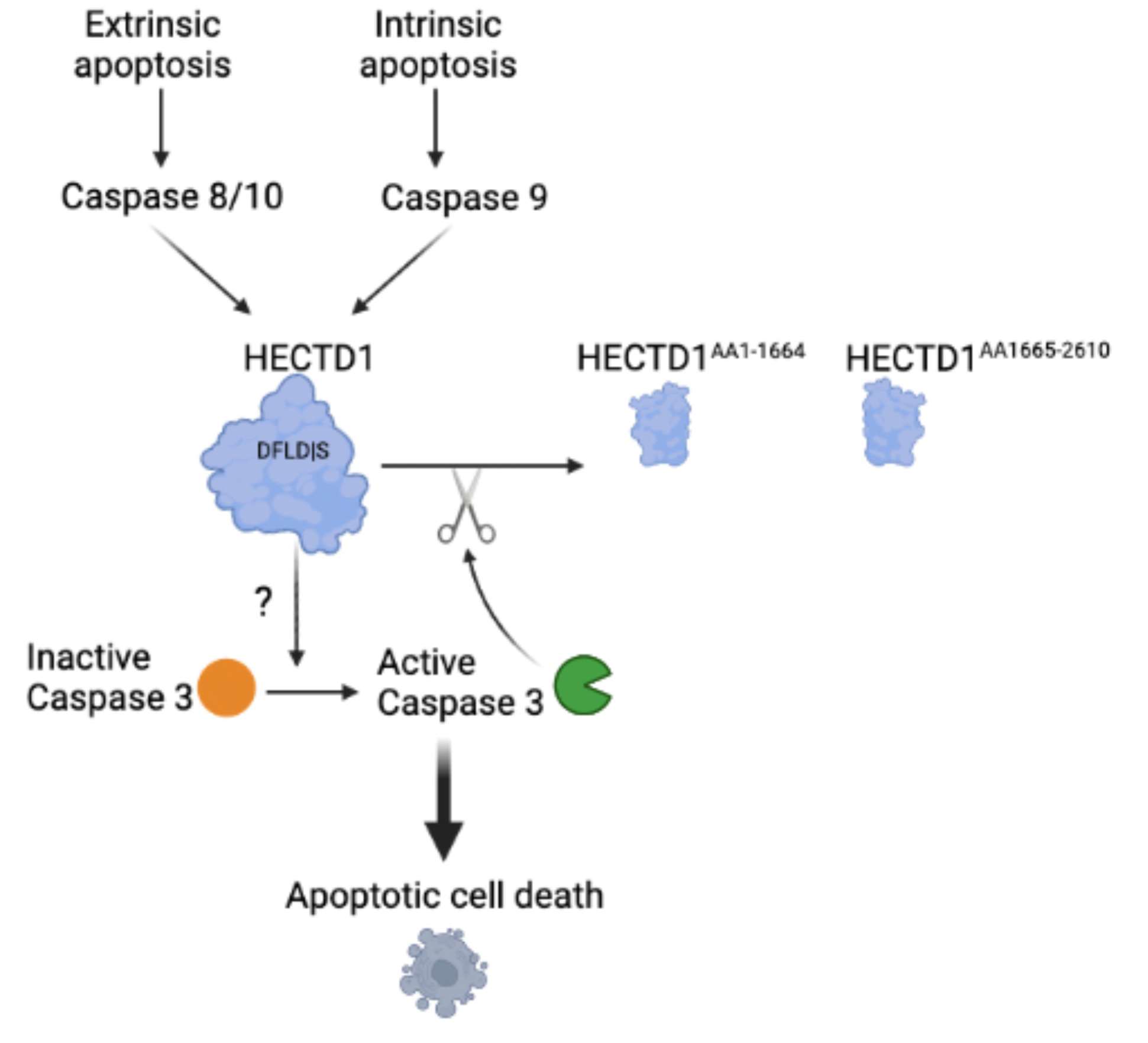
Model for HECTD1-caspase-3 interplay during apoptotic cell death. Our data indicate HECTD1 depletion reduces apoptotic cell death and this is mediated by reduced caspase-3 activity. Therefore HECTD1 is a positive effector of apoptotic cell death. Pulldown experiments using TRABID-NZF as bait revealed that induction of apoptotic cell death leads to an enrichment of ubiquitin signals. Pulldown assay using TRABID NZF suggest ubiquitin chains beyond K63 chains, might be functional during apoptotic cell death, K29 and/or K33- containing ubiquitin linkages since these are preferentially recognised by TRABID. Finally, induction of apoptotic cell death triggers HECTD1 cleavage at Aspartate 1664 and this proteolytic event is mediated by caspase-3 activity. Although HECTD1 C-ter product is as stable as the full length endogenous protein, the function if any of both the N and C-ter products of HECTD1 generated by the caspase cleavage remain to be determined. Created with https://BioRender.com.

The regulation of E3 ubiquitin ligase activity and function by protease is important beyond the realm of apoptotic cell death and caspase activation. For instance, this mode of regulation has emerged as a way for pathogens to interfere with the host immune response and hijack the ubiquitin machinery. The RING E3 RNF20 for instance is cleaved by the viral protease 3CLPro and this inactivates RNF20 ligase activity preventing the degradation of SREBP1, a protein driving lipid metabolism which instead is hijacked by SARS-CoV-2 for viral replication (85). Similarly, cleavage of TRIM7 E3 by an enterovirus encoded 3C protease was recently shown to dampen antiviral immunity (86). Interestingly more than one TRIM7 variant has been identified and these might have different susceptibility to proteolytic cleavage suggesting an evolutionary adaptation yielding variants with distinct antiviral activities in mammals. Whether and how protease-mediated cleavage is used during apoptotic cell death or during the immune response to infection by pathogens to target key components of the ubiquitin system remains poorly understood. However, the evidence that such mechanisms also extend to other components of the ubiquitin system including the ubiquitin like modification ISG15, suggests such mechanisms might be more widespread than originally thought (87). Finally, the discovery of a mutational hotspot in the deubiquitinase USP8 in Cushing’s disease also indicates that proteolytic processing may be an important mechanism targeted during human disease. In this context, somatic mutations in USP8 produce a truncated product with enhanced DUB activity towards EGFR, thus promoting constitutive EGF signalling (88). Future work is needed to further address the role of proteolytic cleavage of ubiquitin components including E3 ubiquitin ligases in the context of caspase-dependent apoptotic and non-apoptotic process and to determine the relevance of these mechanisms in human diseases.

## 6. Author contribution

JDFL formulated the hypothesis, conceived, and managed the project, supervised, and trained students. FS was on the supervisory team and made intellectual contributions. NS carried out the experiments, analysed the data and helped with figure preparation and drafting the manuscript. NS and JDFL wrote the manuscript with help from all the authors.

## 8. Data availability

The datasets analysed during the current study are available in the UNIPROT repository, https://www.uniprot.org/; PROTOMAP https://www.scripps.edu/cravatt/protomap/L

## Abbreviations

HECT: Homologous to the E6AP carboxyl terminus
Poly: (ADP-ribose) polymerase (PARP)
Tumor Necrosis: Factor-Alpha
STS: Staurosporine.

## 10. Acknowledgements

We thank Dr Felix Randow for kindly providing the luciferase NF-κB reporter system.

We also thank Dr Michael Zachariadis (Material and Chemical Characterisation Facility (MC²), University of Bath for his help with the flow cytometry experiments.

