## Supplementary data for "HECTD1 is both a positive regulator and substrate of caspase-3 activity during apoptotic cell death"

**Running Head:** HECTD1 and Apoptosis

**Supplementary information**

**Supplementary Figures**

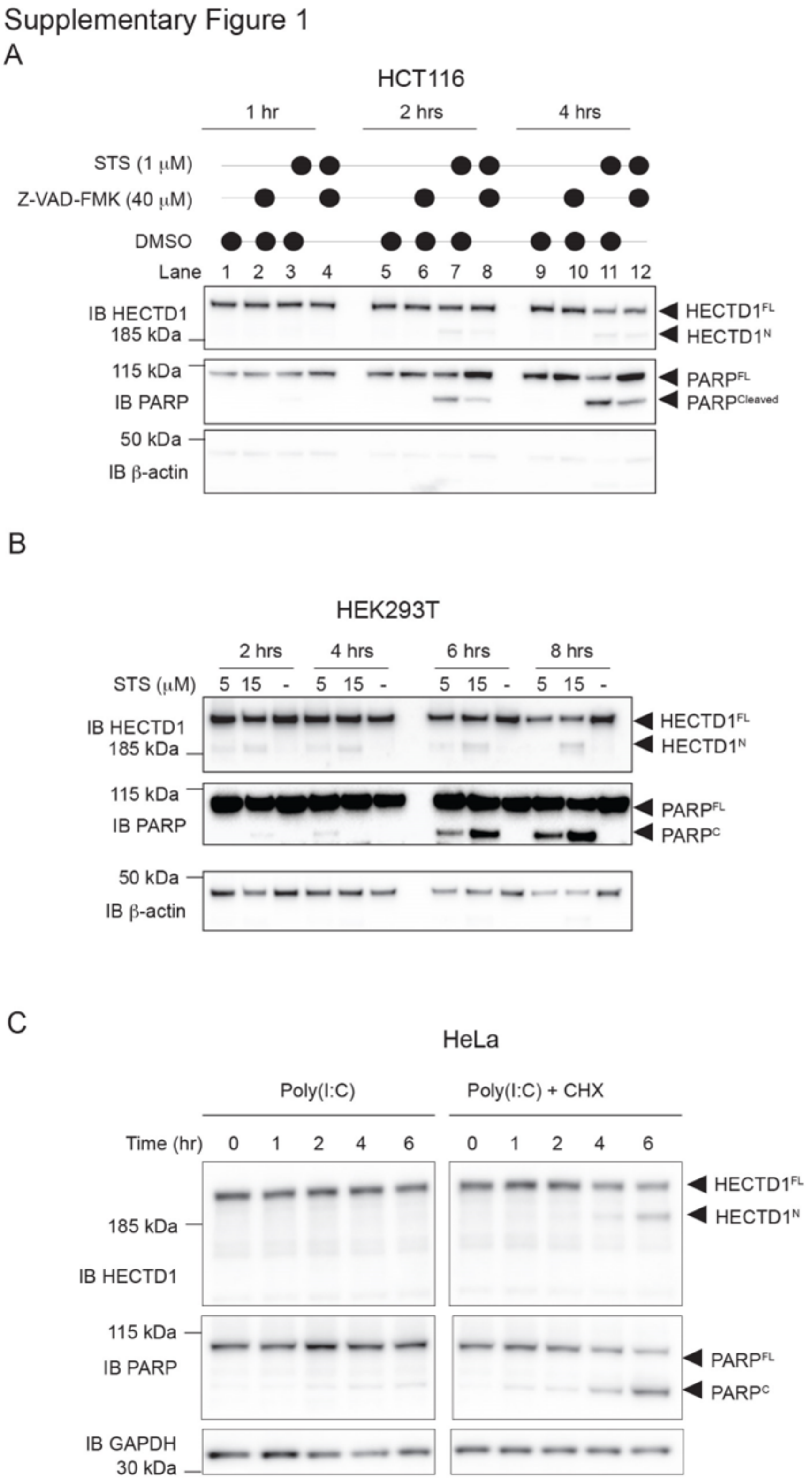


**Supplementary Figure 1. HECTD1 cleavage is detected in HCT116, HEK293T, and HeLa cells. (A)** HCT116 cells were treated with 1 µM STS for 1, 2 or 4 h or and/or were pre-treated with 40 µM z-VAD-FMK for 2 h. Control samples were treated with DMSO. Cells were harvested at the indicated time points, lysed in RIPA, followed by immunoblotting. Samples were loaded on a 4-12% Bis-Tris PAGE and membranes were probed using HECTD1 and PARP antibodies. GAPDH was used as loading control. **(B)** HEK293T cells were treated with the indicated concentrations of STS (5 or 15 μM) or DMSO for the indicated times (2, 4, 6, 8 h) prior to cell lysis. Samples were then handled and analysed by immunoblotting as mentioned in A. **(C)** HeLa cells were treated with poly(I:C) (100 μg/ml) alone or in combination with CHX (10 μg/ml). Cells were harvested and lysed with RIPA at the indicated time points post treatment. HECTD1 and PARP cleavage were monitored by immunoblotting using HECTD1 and PARP antibodies, with GAPDH as loading control. Membranes were cut to enable detection with multiple antibodies.

Supplementary Figure 2

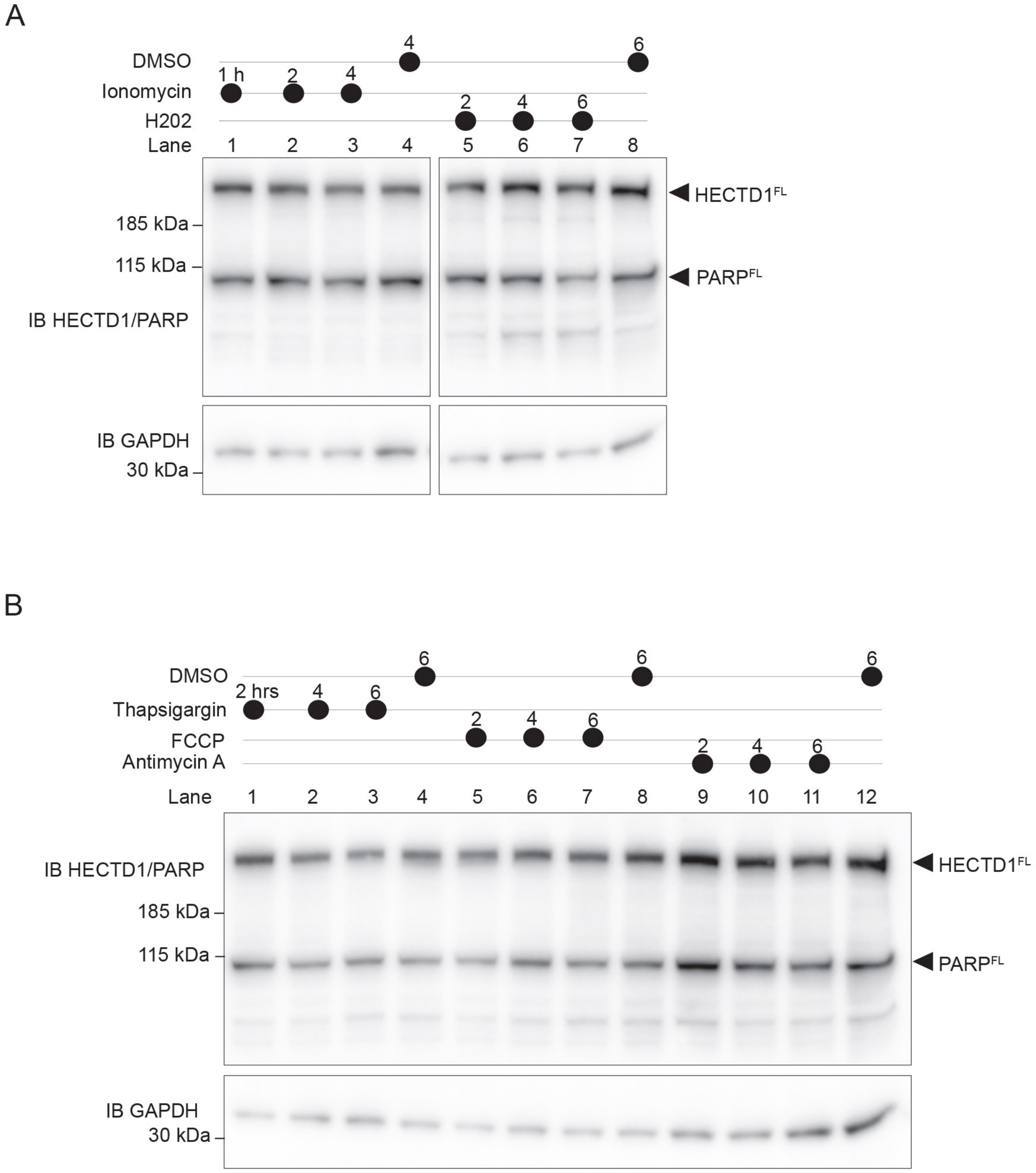


**Supplementary Figure 2. Drugs that induce mitochondrial or ER stress do not induce HECTD1 cleavage in HeLa cells. (A)** HeLa cells were treated with either 3 µM ionomycin for 1 h, 2 h, and 4 h; 250 µM $H_2O_2$ for 2 h, 4 h, and 6 h, or DMSO as control before being lysed in RIPA buffer and analysed by immunoblotting. Samples were loaded on a 4-12% Bis-Tris PAGE and membranes were probed using HECTD1 and PARP antibodies. GAPDH was used as loading control. **(B)** HeLa cells were treated with either 10 µM thapsigargin for 2 h, 4 h, and 6 h; 10 µM FCCP for 2 h, 4 h, and 6 h; 10 µM antimycin A for 2 h, 4 h, and 6 h, or DMSO as control. Immunoblotting was carried out as in A. Membranes were cut to enable detection with multiple antibodies..

Supplementary Figure 3

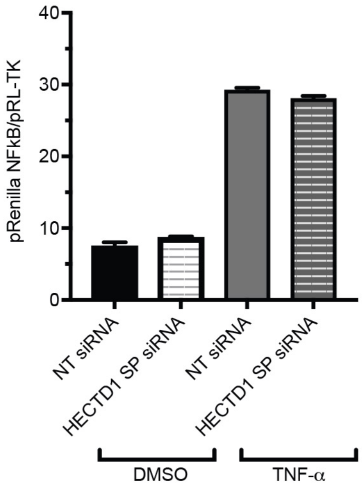


**Supplementary Figure 3. HECTD1 depletion does not affect NF-κB signalling.** The Dual-Luciferase® Reporter Assay (Promega, Madison, WI, USA, E1910) was performed according to manufacturer's instructions. Briefly, HeLa cells were seeded at 8,000 cells/well in a 96-well plate and transfected with either NT or HECTD1 SMARTPool siRNA using FuGENE® HD. 24 h post transfection, cells were transfected with an NF-κB luciferase reporter together with pRL-TK which is used as control (100:1 Firefly:Renilla ratio) for an additional 24 h followed by 6 h treatment with 50 ng/ml TNF-α. Samples were then quantified for Firefly Luciferase and Renilla activity as per the manufacturer's protocol. Firefly to Renilla ratio is shown in a bar graph format. Each condition was performed in triplicate, n = 1 experiment.

Supplementary Figure 4

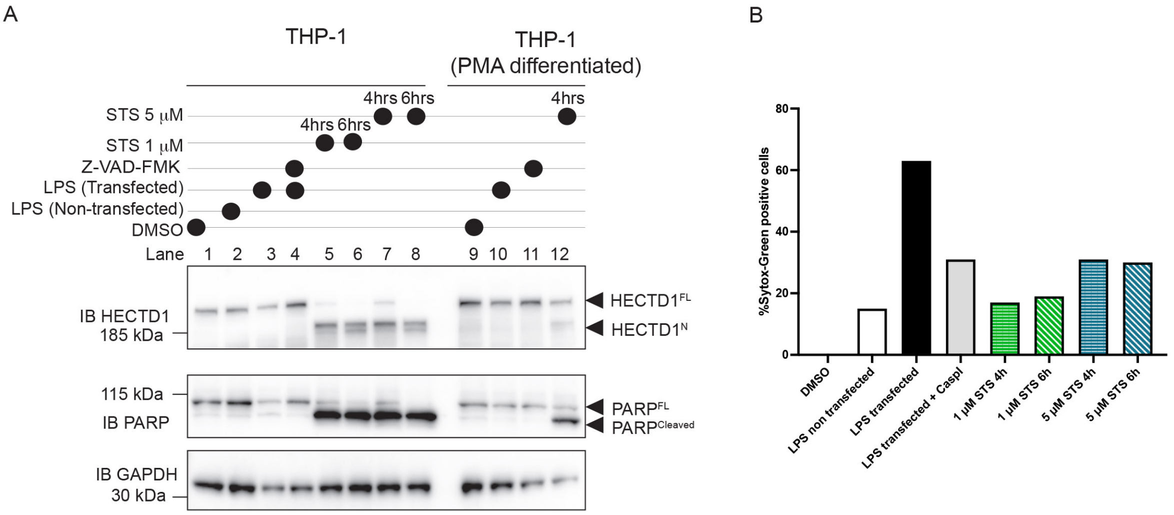


**Supplementary Figure 4. HECTD1 cleavage is not induced by treatments which induce pyroptosis. (A)** The effect of LPS treatment on HECTD1 was assessed in THP-1 and macrophage-like THP-1 cells differentiated using PMA. Cells were treated with 2 μg/ml LPS or transfected with 2 μg/ml LPS using Lipofectamine 2000 for 24 h prior to cell lysis using RIPA buffer. As control HECTD1 cleavage was induced in THP-1 or macrophage-like THP-1 cells using the indicated STS concentration and incubation times. Samples were loaded on a 4-12% Bis-Tris PAGE and membranes were probed using HECTD1 and PARP antibodies. GAPDH was used as loading control. Membranes were cut to enable detection with multiple antibodies. **(B)** Induction of pyroptosis in THP-1 cells was monitored using live-cell Sytox-Green staining (167 nM). Cells were imaged and analysed using Countess 3 Automated Cell Counter prior to processing for Western blot analysis.
